# Lateral gene transfer of anion-conducting channelrhodopsins between green algae and giant viruses

**DOI:** 10.1101/2020.04.15.042127

**Authors:** Andrey Rozenberg, Johannes Oppermann, Jonas Wietek, Rodrigo Gaston Fernandez Lahore, Ruth-Anne Sandaa, Gunnar Bratbak, Peter Hegemann, Oded Béjà

## Abstract

Channelrhodopsins (ChRs) are algal light-gated ion channels widely used as optogenetic tools for manipulating neuronal activity ^1,2^. Four ChR families are currently known. Green algal ^3–5^ and cryptophyte ^6^ cation-conducting ChRs (CCRs), cryptophyte anion-conducting ChRs (ACRs) ^7^, and the MerMAID ChRs ^8^. Here we report the discovery of a new family of phylogenetically distinct ChRs encoded by marine giant viruses and acquired from their unicellular green algal prasinophyte hosts. These previously unknown viral and green algal ChRs act as ACRs when expressed in cultured neuroblastoma-derived cells and are likely involved in behavioral responses to light.

## MAIN

Channelrhodopsins (ChRs) are microbial rhodopsins that directly translate absorbed light into ion fluxes along electrochemical gradients across cellular membranes controlling behavioural light responses in motile algae ^9^. They are widely used in optogenetics to manipulate cellular activity using light ^10^, and therefore there is a constant demand for new types of ChRs with different functions, be it different absorption spectra ^11^, ion selectivity ^12^, or kinetics ^11^. So far, ChRs have only been reported from cultured representatives of two groups of algae: cryptophytes and green algae ^2,13^. Recently, metagenomics proved to be a useful tool to identify novel ChRs, as a new family of anion-conducting ChRs (ACRs) with intensely desensitizing photocurrents were detected in uncultured and yet to be identified marine microorganisms ^8^.

### Channelrhodopsins in metagenomic contigs of putative viral origin

To extend the search for uncharacterized distinct ChRs with potentially new functions, we further screened various metagenomic datasets from *Tara* Oceans ^14–16^. In total, four unique sequences belonging to a previously undescribed family of ChRs were found in five metagenomic contigs from the prokaryotic/girus fractions from tropical and temperate waters of the Atlantic and Pacific Oceans. Two of the contigs were long enough (11 kb and 20 kb) to provide sufficient genomic context (Fig. 1a). We attempted to search for similar sequences in several metagenomic datasets, and located multiple contigs with synteny to the two longer ChR-containing contigs (Fig. 1a, Suppl. File 1). The two contigs recruited two clusters of related fragments (v21821 and v2164382 contig clusters) mostly from marine samples of *Tara* Oceans. Interestingly, the v21821-cluster contigs came from the same South Atlantic station, except for one contig with lower identity and synteny length from a soda lake metagenome ^17^ (LFCJ01000229.1, see Fig. 1a). The v2164382 cluster was more diverse in both the gene order and geography, but all of the recruited contigs came from the marine realm. Although none of those recruited fragments contained ChR genes, they could be utilized in downstream analyses to clarify the origin of the ChR-containing contigs. Surprisingly, inspection of all of those metagenomic contigs demonstrated a typical viral genome organization with intronless genes separated by short spacers and several tRNA genes. The fragments harbored high proportions of genes with affinities to two families of nucleo-cytoplasmic large DNA viruses (NCLDVs), *Mimiviridae* and *Phycodnaviridae*, the two most abundant NCLDV groups in the ocean ^18^, included multiple nucleo-cytoplasmic virus orthologous groups (NCVOGs) ^19^, and demonstrated genome composition similar to these viruses and distinct from the potential host groups (Fig. S1).

**Fig. 1.**
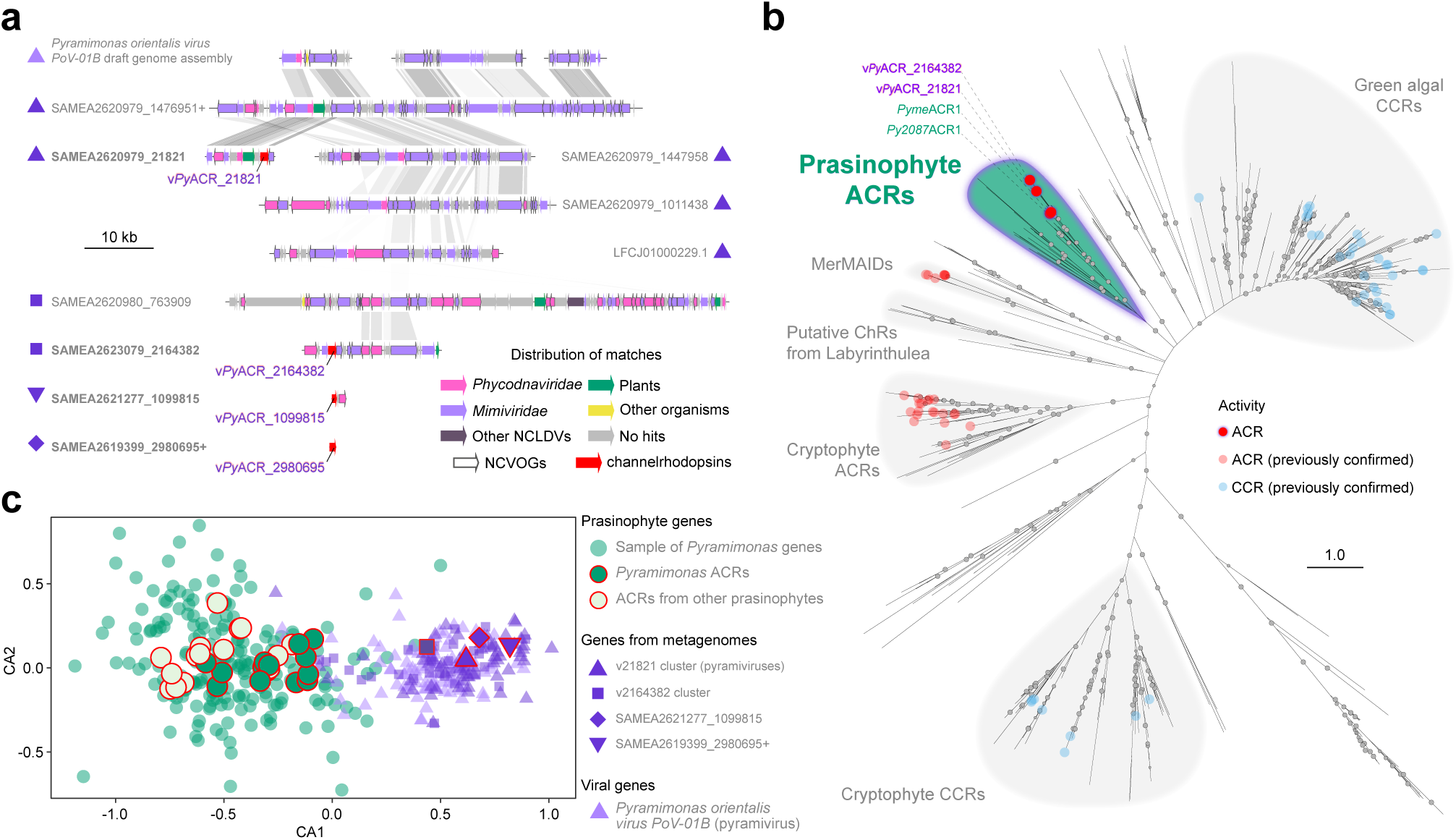
Phylogeny and diversity of viral and green algal ACRs. **(a)** Metagenomic contigs containing ChR genes (in bold) and syntenic contigs used for phylogenetic analyses, alongside corresponding genomic fragments from *Pyramimonas orientalis virus PoV-01B*. Annotated contigs and the draft genome sequence are available in Suppl. Files 1 and 2. **(b)** Unrooted tree showing phylogenetic relationships between confirmed and putative channelrhodopsins including the previously characterized families of green algal CCRs, cryptophyte ACRs and CCRs, MerMAIDs and the prasinophyte/viral ACRs reported here. The four members of the prasinophyte/viral ACR family characterized in the current study are highlighted. The gray circles represent ultrafast bootstrap support values (70-100%), scale bar indicates the average number of amino acid substitutions per site. See Fig. S2 and Suppl. File 3b for the full version of the tree and the alignment. (**c**), Comparison of tetranucleotide composition of ChR genes from prasinophytes and viral metagenomic contigs (encircled in red) against the background of genes from *Pyramimonas* (green), metagenomic contigs (light-blue) and PoV-01B (lilac).

Phylogenetically, the viral channelrhodopsins appeared to be different from the four currently known ChR families and indeed formed a well-supported family of their own (Fig. 1b). Although some members of the viral families *Phycodnaviridae* and *Mimiviridae* are known to harbor other microbial rhodopsins and heliorhodopsins ^20–22^, no virus has previously been described to code for channelrhodopsins. This encouraged us to investigate the function and origins of these ChRs and to identify the corresponding viruses and their putative hosts.

### Homologs of viral ChRs in prasinophyte algae

An initial screen for proteins similar to the viral ChRs yielded homologous genes in several transcriptomes of green algae from two transcriptomic datasets ^23,24^. An in-depth analysis of the available green algal genomes and transcriptomes showed that the homologues of the viral ChRs were strictly confined to three related clades of prasinophytes (paraphyletic assemblage of early diverging unicellular chlorophytes): *Pyramimonadophyceae, Mamiellophyceae* and *Nephroselmidophyceae* with the viral ChRs all clustering together with proteins from the genus *Pyramimonas* (Fig. 1b). Among the previously characterized ChRs, the new family was most similar to green algal CCRs, MerMAIDs and cryptophyte ACRs. The lack of Asp and Glu residues in transmembrane domain 2 (TM2), that are well conserved in green algal CCRs but generally substituted in ACRs (Fig. S3), led us to hypothesize that the putative channelrhodopsins from this new clade might conduct anions, hence the provisional name v*Py*ACRs for “**v**iral ChRs similar to putative ***Py****ramimonas* **ACR**s”.

### Prasinophyte and viral ChRs conduct anions

To examine the function of the new ChR family, we expressed two viral ChRs (v*Py*ACR_21821 and v*Py*ACR_2164382) and one ChR from *Pyramimonas melkonianii* CCMP722 (*Pyme*ACR1) in mouse neuroblastoma x rat neuron hybrid (ND7/23) cells. Two days after transfection, we recorded bidirectional photocurrents under whole-cell voltage-clamp conditions and determined wavelength sensitivity and ion selectivity. While the full-length *Pyme*ACR1 construct expressed well, both full-length viral ChRs showed strong retention in the cytosol and did not yield any photocurrents. We modified the proteins’ N- and C-termini to improve protein folding as well as membrane trafficking and localization (Fig. S4, Methods) ^25^. Even though the viral constructs remained cytotoxic, these modifications enabled us to analyze v*Py*ACR_21821, while for v*Py*ACR_2164382, on the other hand, only singular measurements in standard buffer were possible (Fig. S5).

We determined the action spectra of v*Py*ACR_21821 and *Pyme*ACR1 by recording transient photocurrents upon stimulation with light between 390 and 690 nm (Fig. 2a). v*Py*ACR_21821 is most sensitive (*λ*_max_) to 482 nm light and *Pyme*ACR1 to 505 nm light (Fig. 2a, inset and Fig. S6a). Upon longer illumination, photocurrents of both constructs are non-inactivating during excitation (Fig. 2b,c) but the photocurrent amplitudes of v*Py*ACR_21821 are roughly five times smaller compared to *Pyme*ACR1 (Fig. 2b,c and Fig. S5, S6b). Next, we tested the ion selectivity by recording photocurrents at different membrane voltages and ionic conditions and determined the reversal potential (E_rev_), which is the membrane voltage where inward and outward ion flow cancel each other at a certain ion gradient. Changing the concentration of the conducted ions causes reversal potential shifts (ΔE_rev_). Upon reduction of the external Cl^-^ concentration ([Cl^-^]_ex_) from 150 mM to 80 mM and 10 mM the reversal potential shifts almost equally for *Pyme*ACR1 and v*Py*ACR_21821 (Fig. 2d,e and Fig. S6d,e) to more positive values according to the theoretical Nernst potential (Fig. S6c), whereas it shifts slightly more negative upon replacement of Cl^-^ by Br^-^ or NO_3_^-^, indicating non specific anion conductivity (Fig. 2e and Fig. S6d,e). Replacing external Na^+^ with N-methyl-D-glucamine (NMDG^+^), while keeping [Cl^-^] constant, does not affect the reversal potential and excludes Na^+^ as a transported charge carrier (Fig. 2e and Fig. S6d,e). We therefore conclude that *Pyme*ACR1 and v*Py*ACR_21821 are anion-conducting ChRs (ACRs) that naturally conduct Cl^-^ and potentially Br^-^ and NO ^-^, but do not conduct Na^+^ (Fig. 2e). While for *Pyme*ACR1 the current amplitudes at -60 mV increased at lower [Cl^-^]_ex_, similar to previously described ACRs ^8,26^, in v*Py*ACR_21821 the amplitudes decreased under the same conditions (Fig. 2b,f and Fig. S6f,g).

**Fig. 2.**
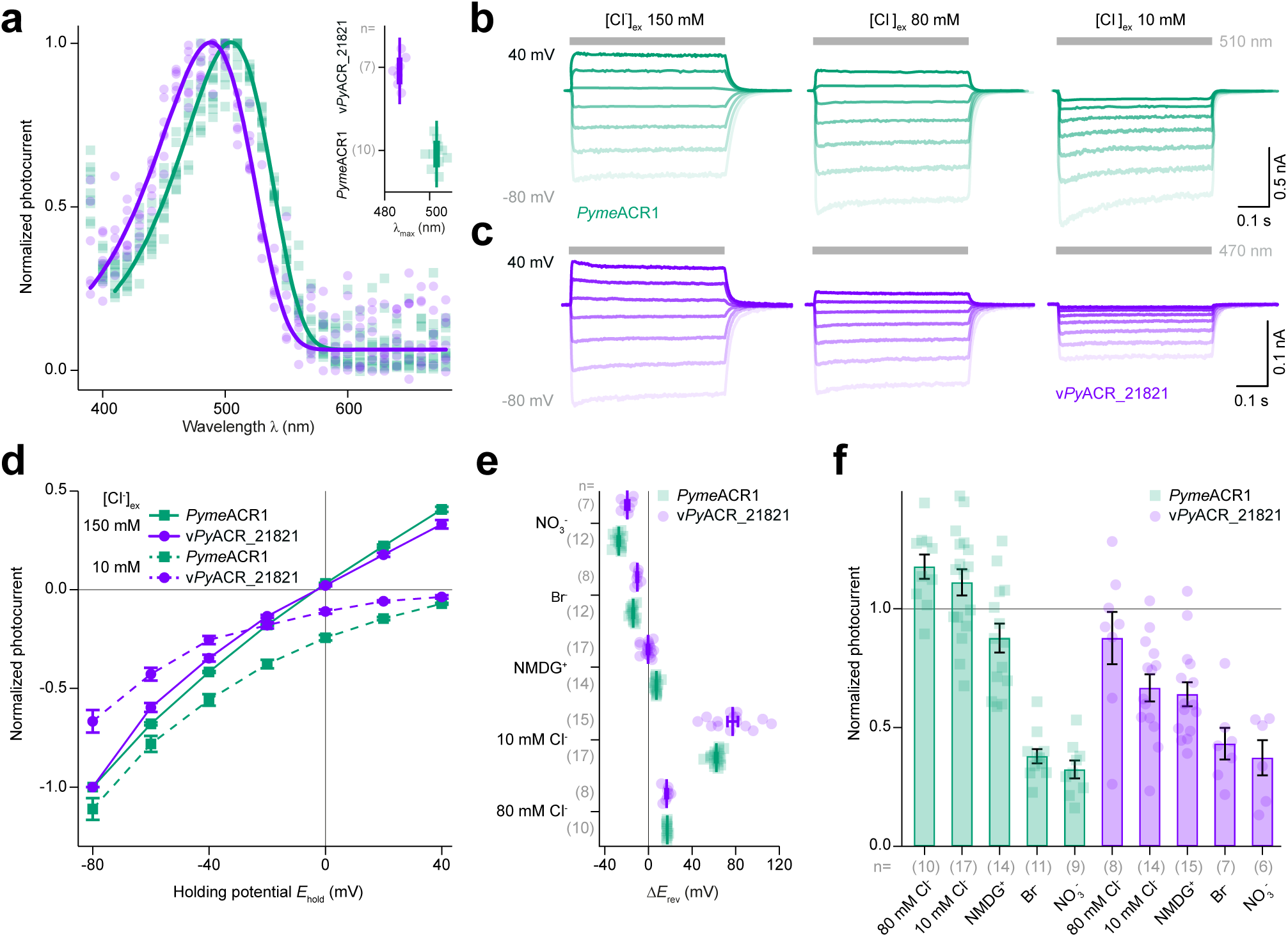
Electrophysiology of viral and green algal ACRs. **(a)**, Action spectra of v*Py*ACR_21821 and *Pyme*ACR1 normalized to the maximum stationary current. Solid lines represent fitted data. Inset shows determined maximum sensitivity (*λ*_max_) for both ChRs. Photocurrent traces of (**b**) *Pyme*ACR1 and (**c**) v*Py*ACR_21821 at indicated extracellular chloride concentrations ([Cl^-^] _ex_) recorded from −80 to +40 mV in 20 mV steps. Gray bars indicate light application of denoted wavelengths. **(d)**, Current-voltage relationship of *Pyme*ACR1 and v*Py*ACR_21821 at [Cl^-^] _ex_ 150 mM (solid line) and 10 mM (dashed line). Photocurrents were normalized to the stationary current at −80 mV with [Cl^-^] 150 mM. **(e)**, Reversal potential shifts (ΔE_rev_) upon exchange of the external buffer. **(f)**, Photocurrent amplitudes at −80 mV upon exchange of the extracellular buffer normalized to the photocurrent amplitude in 150 mM Cl^-^ buffer. Data is shown as single data points (squares or circles), while statistics denote mean ± standard error. The number of conducted experiments (n) is reported in grey. Source data are provided in Suppl. File 5 (a, d-f).

Additionally, we analyzed the photocurrents of a second prasinophyte channelrhodopsin from *Pyramimonas* sp. CCMP2087 (*Py2087*ACR1) (Fig. S7). In contrast to *Pyme*ACR1 and similarly to both v*Py*ACRs, this sequence interestingly has the widely conserved Arg in helix 3 replaced by Gln (position R120 in *Cr*ChR2, see Fig. S3). *Py2087*ACR1 is most sensitive to 509 nm light and the reversal potential of the non-inactivating photocurrents shifts strongly upon reduction of [Cl^-^]_ex_ from 150 mM to 10 mM, indicating that *Py2087*ACR1 is an ACR as well (Fig. S7).

### Prasinophyte ACRs are likely part of the visual system

The fact that green algae appeared to contain previously unknown channelrhodopsins comes as no surprise, since the same group is already known to possess a different family of ChRs, the green algal cation channelrhodopsins (CCRs) ^3,13^. Nevertheless, the distribution of the ACRs is much more narrow as they appear only in some prasinophytes, which is in striking contrast to the CCRs which are distributed widely in chlorophytes and are even present in some streptophytes (Fig. 3 and Fig. S8). At the same time, we notice that in prasinophytes the appearance of CCRs nearly coincides with that of the ACRs. Interestingly, all of the prasinophyte species with genes coding for at least one ChR family have eyespots, the main photosensitive organelle that provides the algae directional sensitivity ^5,27,28^, while for those prasinophyte species that lack the eyespot neither ChR family could be detected. The sister-group relationship between the *Pyramimonadophyceae* and *Mamiellophyceae* ^29,30^ suggests that the last common ancestor of this clade possessed both families of the ChRs, namely CCRs and ACRs, and that they were lost at least three times in this group together with the loss of the eyespot (in the flagellates *Pterosperma* ^31^ and *Micromonas* ^32^ and the cocci *Ostreococcus*-*Bathycoccus* ^33^) (see Fig. 3). These associations indicate that both families of the ChRs are likely involved in sensing light, similar to what is known for CCRs in chlorophyceans ^3,5,13,28,34^.

**Fig. 3.**
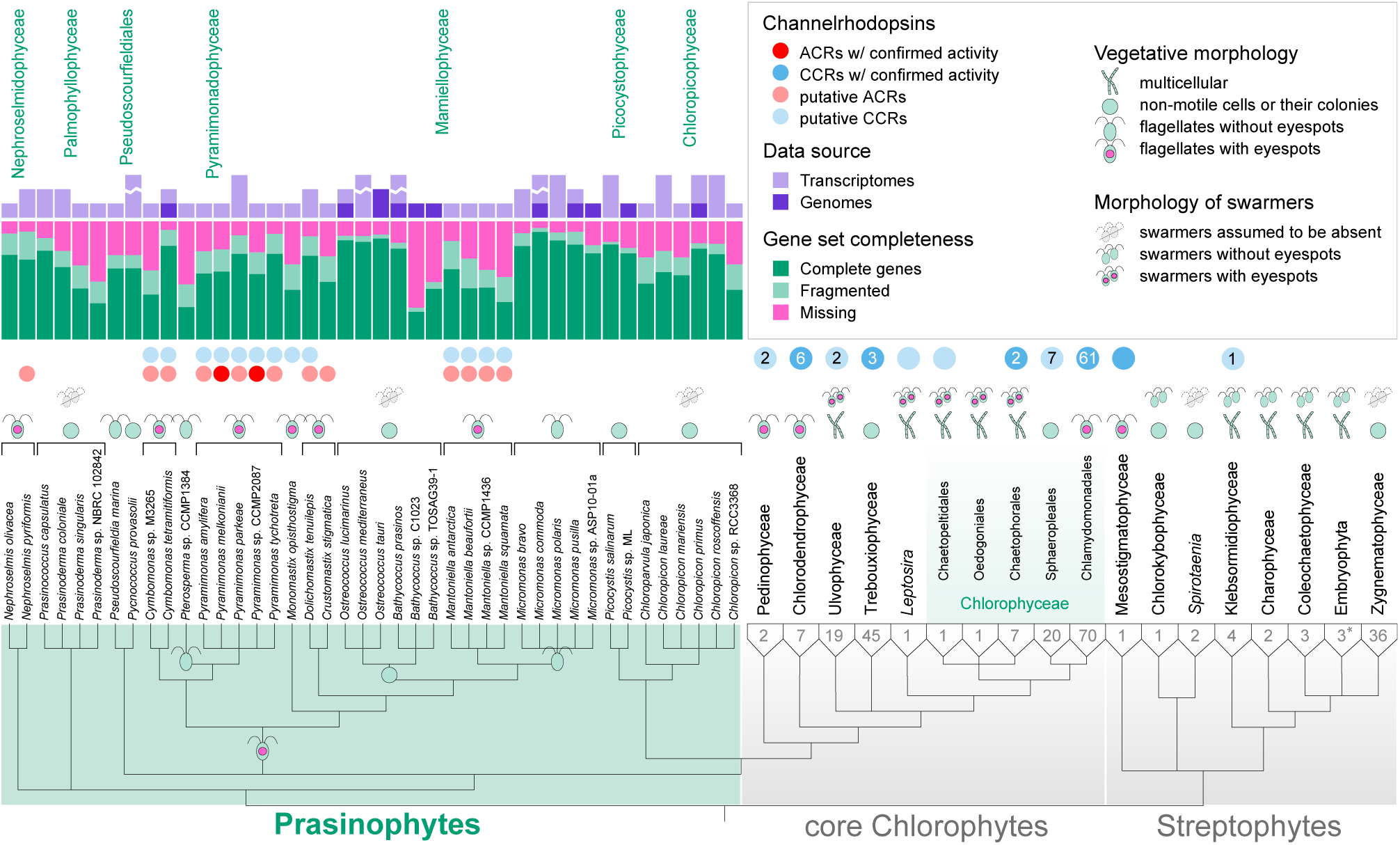
Distribution of ChRs among green algae. Presence/absence of the two families of green algal ChRs in the available transcriptomes and genomes of prasinophytes, as well as summarized counts for the core chlorophyte and streptophyte taxa. For each species several transcriptomes and/or genomes were merged into a single gene set when available. Morphological features with respect to motility and presence of the eyespot are indicated for each prasinophyte species and the dominant types are indicated for the two other groups. The presence of ChRs with confirmed and expected ion selectivities is indicated at the level of species for prasinophytes and at the level of higher taxa for core chlorophytes and streptophytes, with the darker color signifying the presence of at least one ChR with confirmed activity in the corresponding taxon. Numbers indicate the number of all available species with genetic data and the corresponding number of species with ChRs. Note that the embryophytes are represented here by three basal land plant species (*). See a full version of the figure in Fig. S8.

An independent support for the hypothesis of a shared cellular function of the green algal ACRs and CCRs as sensors of light, came from the observation of structural similarities between the two families. Similar to their viral homologs, C-terminally to the rhodopsin domain, full-length prasinophyte ACRs possess a domain with high similarity to response regulators (RR) from two-component regulatory systems (Fig. S9). When searching for remote homology, a similar RR-domain was surprisingly discovered also in green algal CCRs including those coming from prasinophytes, but not in other known groups of ChRs: cryptophyte ACRs and CCRs and the MerMAIDs (see Fig. S9a). In green algal CCRs, this domain corresponds to one of the three previously noticed conserved regions, con2, in the C-terminal extensions of chlorophycean ChRs ^34^. An RR-like domain could also be identified in putative channelrhodopsins from Labyrinthulea, a marine heterotrophic stramenopile group (see Fig. S9). This indicates that this domain is not restricted to ChRs from green algae and was likely present in the last common ancestor of at least the two families of green algal ChRs and their homologs from Labyrinthulea. Interestingly, this domain organization is reminiscent of His-kinase rhodopsins (HKRs), a group of enzymerhodopsins from green algae exemplified by COP5–COP12 from *Chlamydomonas reinhardtii* ^5,35,36^. Yet, in these proteins the rhodopsin and the RR domains are invariably associated with a corresponding transducer (His-kinase) domain as part of a complete two-component system typical of other eukaryotic sensor systems ^37,38^ (see Fig. S9). In addition, in contrast to the classical response regulators, including the RR domain of the enzymerhodopsins, the Asp-57 residue that serves as the phosphorylation site as well as the highly conserved residues Ser/Thr-87 and Lys-109 that are responsible for the conformation change mediated by phosphorylation ^39,40^ are mutated in nearly all of the cases in the ChR sequences from green algae, viruses and Labyrinthulea (see Fig. S9b). Asp-less receiver domains (ALDs) are known and are widespread across the tree of life ^41^. Asn-57 at the phosphorylation site, as seen in the prasinophyte and viral ACRs and the putative labyrinthulean ChRs, is the most frequent substitution in ALDs and has the potential to undergo deamidation to aspartate ^41,42^. At the same time, substitution of the highly conserved Ser/Thr-87 and Lys-109 residues puts the RR-like domains of the ChRs, including those of the prasinophyte ACRs, in a minority position even among ALDs. This likely indicates that the RR-like domains in ChRs do not undergo phosphorylation and conformational changes and thus function as constitutive signals ^41^ or lack a signal transduction function altogether.

The C-terminal extension of *C. reinhardtii* ChR1 is known to participate in regulation and trafficking ^35,43^. Moreover, there is evidence that in *Chlamydomonas*, ChRs and HKRs have similar spatial distribution on the cellular membrane ^36,37^ and that at least two proteins from these families, ChR1 and COP8 utilize intraflagellar transport (IFT) for their delivery to the eyespot and flagella ^36^. We thus tentatively hypothesize that the RR and RR-like domains of HKRs and ChRs, respectively, might function as a shared trafficking signal, in addition to their signaling function or even instead of it, in the case of the ChRs. Note in this context the lack of such domains in both, the cryptophyte ChRs (see Fig. S9a) and the cryptophyte sensory rhodopsins ^44^. It must be noted that since the close association of the eyespot with respect to microtubular roots is characteristic only to the UTC clade (Chlorophyceae, Trebouxiophyceae and Uvlophyceae) and is rarely observed in other chlorophytes (e.g. in *Nephroselmis*) ^45,46^, the trafficking signal is not expected to be directly associated with IFT *per se*.

### *v*Py*ACRs indeed come from giant viruses*

With a plausible hypothesis about the function of the prasinophyte ACRs in hand, we turned to the analysis of the origin of their putative viral homologues that motivated our study in the first place. None of the available NCLDV genomes demonstrated close similarity to the metagenomic fragments in terms of synteny and sequence identity, including those viruses that are known to infect green algae ^47^ and specifically prasinophytes ^48^. First we noticed that the v*Py*ACRs originated from a particular clade of prasinophyte ACRs from *Pyramimonas* as suggested by phylogenetic analysis (see Fig. S2) and that members of the same genus are the only green algal group with ACRs to be reported to demonstrate viral infections in natural populations and in culture ^49,50^ (see also Fig. S10). We thus focused on the sole isolate of a *Pyramimonas*-infecting virus: *Pyramimonas orientalis virus PoV-01B* (*Mimiviridae*) isolated in Norway two decades ago ^50^. Although since then the virus has been lost in culture and no complete genome was released, enough genetic data had been generated to assemble a draft genome. It appeared that the PoV-01B genome assembly indeed contained large fragments syntenic with contigs from the v21821 cluster (see Fig. 1a), which provided a direct connection between the metagenomic contigs and a described viral isolate. Interestingly, despite the synteny, no other sequence from this cluster besides SAMEA2620979_21821 itself, contained the channelrhodopsin gene at the corresponding locus, and the same applied to the PoV-01B genome which did not contain any rhodopsin genes in the sequenced parts of the genome either. Analogously, the metagenomic contig with the highest similarity to SAMEA2620979_21821 which came from the same sampling location showed exactly the same gene arrangement with the exception of the lack of the channelrhodopsin gene (see Fig. 1a). That the ChR-containing viruses from this cluster are indeed relatively scarce was further supported by the fact that even at that particular station they were outnumbered by their ChR-lacking counterparts (Fig. S11). This diversity in gene arrangement and the fact that v*Py*ACR homologs were found in green algae warranted us to test whether the ChR-containing contigs could actually come not from independent viruses, but from fossilized viral fragments in prasinophyte genomes. We compared tetranucleotide composition for the viral ChRs and their green algal homologs against the background of other genes from the metagenomic contigs and PoV-01B on one hand and from the algae on the other. The two resulting clouds of genes were generally well separated and the viral ChR genes fell separately from their algal homologs and well within the viral cloud, thus rejecting the hypothesis that the ChR genes are not part of the original viral genomes (Fig. 1c).

Notwithstanding the monophyly of the viral ChRs, the viruses from which they came appeared to belong to at least two different lineages. First, we noticed that the PoV-01B genome showed synteny with only one of the two clusters of metagenomic contigs, and furthermore that the two clusters showed virtually no overlap in gene composition and had different proportions of mimivirid and phycodnavirid genes (see Fig. 1c). That the two clusters might represent two viral lineages and not merely two disjoint genomic locations, was hinted at by phylogenetic analysis of the sole shared gene, the D5-like helicase/primase, which placed contigs from the v21821 cluster together with one of the two helicase/primase genes from PoV-01B confidently among mesomimiviruses (*Mimiviridae*) and those from the v2164382 cluster in a separate but related clade (Fig. S12). Analogously, based on gene composition analyses (Fig. S13), the longest contigs from the v21821 cluster, along with PoV-01B could be clearly attributed to mesomimiviruses. In these analyses, the sole long contig from the second cluster showed more affinity to the *Phycodnaviridae*, and in particular to *Raphidovirus*, a virus with a unique position among the members of the family ^51^. Finally, phylogenetic analysis of highly conserved genes placed PoV-01B well among the v21821-cluster contigs and the whole clade, again within the mesomimiviruses (Fig. S14). The only non-marine representative of this cluster, the contig LFCJ01000229.1 from a soda lake, appeared as the basalmost and early branching member of this clade. The long v2164382-cluster contig was resolved as a deep branching member of the *Phycodnaviridae*, with *Raphidovirus* as the closest cultured virus, although the exact branching order remained unclear. The putative viral genomes coding for the two other ChRs remain unidentified.

### *The v*Py*ACR-containing viruses infect* Pyramimonas *algae*

Several lines of evidence suggest that the ChR-harboring viruses and at least some of their relatives infect prasinophytes from the genus *Pyramimonas*. First of all, as noted above the shared origin of all four viral ACRs could be traced back to this particular prasinophyte group. Noteworthy, they branch within the clade composed of ChRs from species of the monophyletic subgenus *Vestigifera* and thus these algae can be conclusively identified as the donors of the viral ChRs (Fig. 1b, Fig. S2 and Fig. S15). Note that the host of PoV-01B, a member of the v21821-cluster, is a *Pyramimonas* species from the same clade (see Fig. S15). Moreover, among the genes associated with the two putative viral lineages, four, including a gene for plastidic ATP/ADP-transporter, were found to have no homologs in other known viral genomes, but instead were detected in plants (see Fig. 1a). The three genes that were distributed widely among green algae demonstrated the highest similarity specifically to corresponding homologs from *Pyramimonas* (see Fig. S8). In this respect, the viruses from which the sole non-marine fragment (LFCJ01000229.1, most distant to the ChR-containing contig SAMEA2620979_21821) analyzed here comes from, must have a different host as no *Pyramimonas* species are known from soda lakes.

## CONCLUSIONS

Here we provide characterization of a new family of anion-conducting ChRs with intriguing physiological and ecological implications. Although we identified the first members of the family as putative viral proteins, these ChRs were found to be widespread among prasinophyte green algae. In motile members of this group we find both, ACRs and close relatives of CCRs from other green algae, and furthermore similarities in the C-termini in proteins from the two families and evolutionary association with the eyespot imply that both, ACRs and CCRs, are utilized by motile prasinophytes for light sensing. It remains to be discovered how ACRs and CCRs co-operate in regulating motility in these algae and what allows other green algae to rely solely on CCRs apparently without concomitant simplification in swimming behavior (compare e.g. the behavioral spectra of *Pyramimonas* ^52^ and *Chlamydomonas* ^28^).

The viral homologs of the prasinophyte ACRs that initially prompted our study, represent a relatively recent acquisition from host genomes by *Pyramimonas*-infecting viruses as it does not predate the diversification of the genus into morphologically distinct lineages, yet nucleotide sequences of the corresponding genes lost trace of their algal origin. Despite coming from at least two viral lineages, the mimivirids pyramiviruses and a putative phycodnavirid clade, the four distinct ChRs from viruses form a monophylum and thus originate from a single alga-virus lateral gene transfer. The question suggests itself: Why would viruses carry channelrhodopsin genes, what selective advantage do they provide? Given the likely role of their algal homologs in sensing light and the preservation of the intracellular C-terminus in viral ACRs with respect to host ChRs, we propose the hypothesis that the role of the viral ACRs is manipulation of host’s swimming behavior. A similar hypothesis has been proposed for the Group-I and Group-II viral rhodopsins, a different family of microbial rhodopsins found in genomes of several mesomimiviruses ^20,21^, that are hypothesized to function as pumps or channels, respectively ^21,53^. The benefits of modifying phototactic or photophobic responses of the host cell by the virus might range from avoidance of oxidative stress to optimization of photosynthesis for the needs of the virus.

## Supporting information

Submitted to Genbank - Constructs

## Acknowledgments

We thank José Flores-Uribe for help with bioinformatics, Eunsoo Kim for providing us an unpublished transcriptome assembly of *Cymbomonas tetramitiformis*, Stuart D. Sym and Charles J. O’Kelly for their comments on *Pyramimonas* taxonomy and Richard Pienaar and Stuart D. Sym for providing us with micrographs of viral infection in *Pyramimonas pseudoparkeae*. This work was supported by Israel Science Foundation grant 143/2018 (O.B.), the Milgrom Foundation (O.B.), Research Council of Norway project VirVar 294363 (R.-A.S.), and German Research Foundation grant SFB 1078 B2 (P.H.). P.H. is a Hertie Senior Professor for Neuroscience supported by the Hertie Foundation. O.B. holds the Louis and Lyra Richmond Chair in Life Sciences.

## Author contributions

O.B. and J.W. conceived the project. A.R. and O.B. performed bioinformatic analyses. P.H. and J.O. designed molecular characterization. J.O. and R.G.F.L acquired and analyzed electrophysiology and imaging data, respectively. R.-A.S. and G.B. performed genome sequencing of the PoV-01B virus. A.R., J.O., and O.B. wrote the paper, with contributions from all authors.

## Supplementary information

### Materials & Methods

#### Metagenomic contigs

The viral ChRs were found in five assembled contigs from *Tara* Oceans: SAMEA2620979_21821, SAMEA2623079_2164382, SAMEA2621277_1099815, SAMEA2619548_2902552 and SAMEA2619399_2980695 (the SAMEA* prefixes refer to corresponding NCBI biosamples). Although the search was performed on several available assemblies of *Tara* Oceans, all five contigs come from the assembly generated previously by ^1^. The two shortest contigs contained two overlapping fragments of a single ORF and could be merged together thanks to an identical overlap of 225 bp. ORFs were annotated using GeneMarkS v. 4.32 ^2^ in the eukaryotic viral mode and prokka v. 1.14.5 ^3^ in the viral mode (giving preference to the GeneMarkS gene boundaries in cases of conflict) with manual corrections and tRNA genes were annotated using tRNAscan-SE v. 2.0.3 ^4^. To increase the set of genes suitable for identification of the corresponding ChR-containing contigs additional longer contigs were recruited by searching for contigs containing homologs to at least four genes from the ChR-containing contigs using blastp v. 2.2.31+ ^5^. One of the longest recruited contigs without ChRs, SAMEA2620979_1476951 (34,130 bp) could be significantly extended further by stitching it with a different contig retrieved from the same marine station, SAMEA2620979_1432764 (39,065), thanks to an exceptionally long overlap of 11,902-11,903 bp with an identity level of 88.5%.

*Draft genome assembly of* Pyramimonas orientalis virus PoV-01B. Isolation and culture of *Pyramimonas orientalis virus PoV-01B* were described previously ^6^. Shotgun libraries of randomly sheared, end-repaired DNA from PoV-01B (1-2 Kb) were prepared by Lucigen (https://www.lucigen.com/) using the pSMART-HCKan cloning vector (Lucigen,WI, USA). Clones were sequenced by Sanger sequencing using the MegaBACE 1000 and 4000 instruments (Symbio Corporation, CA, USA) for a total yield of 9666 reads. Base-calling was performed with phred v. 0.020425.c ^7^. The data was assembled with phrap v. 0.990329 ^8^ resulting in 141 contigs (total length 689,147 bp, N50 = 12,634 bp, L50 = 15). For further analysis the contigs were trimmed at low-quality regions and those longer than 2000 bp were taken for downstream analysis (except for Contig39 which contained the gene for helicase D10). Additionally, a Nextera library was prepared from an infection experiment and a pilot MiSeq run was performed with a total yield of 9225 reads mappable to the genome draft. The contigs were checked for potential overlaps undetected by the assembler due to sequencing errors with blastn and manually joined with the assistance of the Illumina data when needed. The resulting assembly amounted 36 contigs (total length 523,791 bp, N50 = 23,083 bp, L50 = 7, raw read alignment rate 91.12%, coverage 12.64 reads/bp). Annotation was performed using the same pipeline as for the metagenomic contigs. The genetic data was initially intended to assist discovery of phylogenetic markers ^6,9^ and was not planned to be released as a genome assembly because of incompleteness and sequencing errors. Nevertheless, we managed to retrieve 11 out of 12 highly conserved genes (see Fig. S14), and thus the assembly might be considered close to complete. With a notice of remaining sequencing errors, the assembly is released here as a supplement file (Suppl. File 2), the sequences of phylogenetic markers in Suppl. File 6a.

The search for potential rhodopsin genes was performed with blastp and tblastn searches against assembled contigs and raw reads.

#### Green algal ChRs

Two transcriptomic datasets were recruited in the search for viral ChR homologs: MMETSP ^10^ re-assemblies (https://doi.org/10.5281/zenodo.746048) and 1KP ^11^ assemblies. After green algae appeared as the only group containing homologs of the viral ChRs, additional genome and transcriptome assemblies from green algae from NCBI and JGI were added to the dataset. Previously unannotated genomes were annotated with GeneMark-ES v. 4.38 ^2^ in the self-training mode with default settings. For genome assemblies with low N50 values, the gene annotation was performed by running training with the minimum contig length lowered to 5000 (NCBI assembly GCA_004000685.1) or by using models trained on closely related genomes (NCBI assemblies GCA_001630525.1 [GCA_002588565.1 as reference], GCA_002317545.1 [GCA_004335915.1], GCA_003612995.1 [GCA_002897115.1], GCA_003613005.1 [GCA_002897115.1], GCA_004335885.1 [GCA_002284615.1], GCA_004335895.1 [GCA_001662425.1], GCA_004764505.1 [GCA_004335915.1], GCA_008037345.1 [GCA_002814315.1]). The transcriptome of *Pyramimonas tychotreta* was obtained by clustering contigs from kmers assemblies provided in NCBI SRA for runs SRR4293310-SRR4293315, SRR4293322 and SRR4293323 (Bioproject PRJNA342459) using CD-HIT v. 4.6 ^12^ at the identity level of 99%. Coding sequences for all of the transcriptomes were predicted with TransDecoder v. 5.5.0 (https://github.com/TransDecoder/TransDecoder). Channelrhodopsins were searched for by running a custom pipeline combining pfam profile matching assisted by hmmer v. 3.2 (http://hmmer.org/) with NCBI blast ^5^ searches and confirming their identity by alignment and phylogenetic reconstruction (see below).

Species assignments of some algal strains were corrected or updated as indicated in Suppl. Table 2 with the biggest changes affecting the recently revised genera *Picochloron* and *Micromonas* ^13,14^ Green algal transcriptome and genome assemblies were combined at the level of species, their redundancy was reduced by clustering protein sequences with CD-HIT v. 4.6 at the identity level of 99%. The completeness of the resulting per-species gene sets was tested with BUSCO v. 4.0.2 ^15^ using the viridiplantae_odb10 reference dataset.

Prasinophyte ACRs were defined as those of taxonomically and structurally close ChR sequences including the four confirmed ACRs. Green algal CCRs were defined as those ChRs from green algae which fall within the smallest clade encompassing all known green algal ChRs with confirmed CCR activity (see Fig. S2). Although no experimental evidence exists for the cation-conducting activity of the prasinophyte proteins falling within this clade, primarily because of their cytotoxicity (pers. observations; two of these proteins were unsuccessfully tested in ^16^), they nevertheless possess the well-conserved Asp positions of the green algal CCRs (see Fig. S3). This definition effectively excluded several green algal putative ChRs of uncertain activity (see Fig. S2) which nevertheless had little influence on the picture of the overall distribution of CCRs as most of those proteins came from species also containing proteins from the defined CCR clade.

Phylogenetic relationships between green algae were adopted from ^17^ and further refined based on ^13^. Two cases of uncertain phylogenetic position were verified by extracting and blasting rbcL and 18S sequences: *Scourfieldia* sp. M0560/2 (1KP assembly EGNB, related to *Tetraselmis* and *Scherffelia*) and Trebouxiophyceae sp. KSI-1 (NCBI assembly GCA_003568905.1, belongs to the *Watanabea* clade). Morphological descriptions and habitats were taken from AlgaeBase (https://www.algaebase.org) and primary literature. Vegetative stages were assigned to one of the following categories: 1) multicellular thalli (filamentous, parenchymatous or pseudoparenchymatous); 2) cocci or colonies/clusters of non-motile cells; 3) flagellates a) with or b) without eyespots. Algae with life-cycles involving alternating vegetative flagellated and non-motile resting phases were coded as flagellates. The algae from the first two categories were further supplied with the information about the presence of non-vegetative flagellated stages (zoospores and/or gametes): 1) those that are assumed to have no flagellated stages; 2) those that have at least one flagellated stage, a) with or b) without eyespots in at least one such stage. Only direct morphological evidence was taken into consideration for species or genera (whenever the corresponding characteristic was included in the generic diagnosis), except for Zygnematophyceae which are known to lack motile stages as a clade ^18^. The complete list of analyzed transcriptomes and genomes is provided in Suppl. File 4.

Phylogenetic relationships between the *Pyramimonas* species from which transcriptomes were available were analyzed by extracting nucleotide sequences of ribulose-1,5-bisphosphate calboxylase/oxygenase large subunit (rbcL) gene using blast from the corresponding datasets and recruiting previously published sequences ^19,20^. Sequences from *Cymbomonas* were included for outgroup rooting. Alignments were obtained with mafft and analyzed with iqtree (automatic model selection) without trimming.

#### Analysis of ChR domain organization

Initial analysis of ChR domain organization was performed by running the InterProScan v. 5.36-75.0 pipeline ^21^ on individual sequences. This strategy allowed the identification of a conserved region in the C-terminal extensions of prasinophyte ACRs as a response regulator domain (RR), but failed to identify the con2 region (see ^22^) in green algal CCRs that could be aligned with it. To have an independent confirmation of this homologization, different sets of ChRs were created based on well-defined phylogenetic clades and aligned using mafft (G-INS-i). The alignments were converted into protein profiles with HHmake from HH-suite v. 3.2.0 ^23^ (requiring 50% coverage to record a match), supplied with secondary structure predictions (addss.pl) and analyzed with HHsearch against the pdb70 v. 200101 database. The final alignment was created from complete sequences of prasinophyte and viral ACRs, green algal CCRs, green algal HKRs and CheY as the reference with mafft (G-INS-i). Neither InterProScan, nor HHsearch or alignment could identify any domain with homology to RRs in the C-termini of cryptophyte ACRs and CCRs and MerMAIDs.

#### Tetranucleotide composition analysis

Two separate analyses of tetranucleotide composition were performed: (1) whole-genome composition was calculated for both strands for metagenomic contigs, viruses from the *Phycodnaviridae* and *Mimiviridae*, as well as representatives of their photosynthetic host groups (stramenopiles, green algae and haptophytes); and (2) individual gene’s composition was calculated for CDS sequences of the ACR genes from the metagenomic contigs and prasinophytes. For the second analysis, random samples of non-ChR genes with CDSs at least 200 bp were taken as background: 100 genes from metagenomic contigs and PoV-01B, each, and 40 genes from each of the following *Pyramimonas* transcriptomes: MMETSP0059, MMETSP1081, MMETSP1169, MMETSP1445 and PRJNA342459-36897 (*P. tychotreta*). The tetranucleotide compositions were analyzed with correspondence analysis (cca function from the vegan v. 2.5-5 ^24^ package).

#### Phylogenetic analysis

The alignment of the channelrhodopsin sequences for phylogenetic analysis was performed as follows. All of the collected putative channelrhodopsin protein sequences from the transcriptomic and genomic datasets as well as reference sequences were clustered with CD-HIT at the identity level of 98% and aligned with mafft (G-INS-i). The rhodopsin domain was extracted from the alignment and the positions occupied by gaps in less than 50% of the sequences were trimmed. The resulting alignment was clustered at 100% identity and used to perform phylogenetic analysis with iqtree v. 1.6.10 ^25^ (automatic model selection, 1000 ultrafast bootstrap replicates).

The phylogenetic relationships within the families *Phycodnaviridae* and *Mimiviridae* and the metagenomic contigs were resolved as follows. Homologous genes were collected with GET_HOMOLOGUES v. 11042019 ^26^ using all three available algorithms (BDBH, COG and OMCL with inflation values 1, 1.5, 2, 3, 4 and 5) with an e-value threshold of 1e-3 for full genomes of cultured viruses (excluding the known phycodnavirid outliers *Medusavirus, Mollivurus* and *Pandoravirus*) and the resulting clusters were filtered by requiring no paralogs and a taxonomic coverage of greater than 90%. Homologous genes from the rest of the genomes and metagenomic contigs were fetched by taking best hits with diamond v. 0.9.24 ^27^ (e-value threshold of 1e-5 and subject coverage of at least 50%). The homologs were aligned using mafft v. 7.310 ^28^ (G-INS-i method), trimmed with trimAl v. 1.4.rev15 ^29^ (the “automated1” mode) and the phylogeny was reconstructed with iqtree specifying the orthologs as individual partitions, picking optimal partitioning scheme and substitution models and testing the resulting ML phylogeny with 1000 ultrafast bootstrap replicates.

The D5-like helicase/primase dataset was created by blasting the protein sequences of the helicase-primase genes from the metagenomic contigs against NCLDV protein sequences. The sequences were aligned with mafft (G-INS-i), trimmed with trimAl (automated1) and the phylogeny was reconstructed with iqtree (selecting best-fit model, applying 1000 ultrafast bootstrap replicates). To increase the resolution the process was repeated by focusing on the clade covering the genes from the metagenomic contigs.

#### Gene sharing analyses

For the gene sharing analyses, orthogroups for the longest metagenomic contigs and *Phycodnaviridae* and *Mimiviridae* genomes were collected with GET_HOMOLOGUES (COG algorithm) with an e-value threshold of 1e-3. Genome clustering (Ward’s method) was performed on the genome-by-genome matrix of relative numbers of shared orthogroups. Genome ordination was obtained with correspondence analysis run on the genome-by-orthogroup presence-absence matrix.

#### Molecular biology

For electrophysiological recordings in ND7/23 cells, human/mouse codon-optimized sequences encoding v*Py*ACR_21821, v*Py*ACR_2164382, *Pyme*ACR1, and *Py*2087ACR1 were synthesized (GenScript, Piscataway, NJ) and cloned in frame with mCherry into the pmCherry-C1 vector using *Nhe*I and *Age*I restriction sites (FastDigest, Thermo Fisher Scientific, Waltham, MA). To improve the membrane localization, v*Py*ACR_21821 and v*Py*ACR_2164382 were further subcloned in frame with eYFP into the pEYFP-N1 vector, using Gibson assembly ^30^. As previously reported ^31^, a membrane trafficking sequence (KSRITSEGEYIPLDQIDINV) and an endoplasmic reticulum release sequence (FCYENEV) flanked the fluorophore eYFP, and the N-terminus was extended (MDYGGALSAVGLFQTSYTLENNGSVICIPNNGQCFCLAWLKSNG). Furthermore, the last 131 amino acids of the C-terminus were truncated.

Molecular cloning was planned using NEBuilder v. 2.1.0 (New England Biolabs Inc., Ipswich, MA) and SnapGene v. 4.3+ (GSL Biotech LLC, Chicago, IL).

The sequences of the codon-optimized CDSs of the prasinophyte and viral ACRs and the corresponding translations were deposited in Genbank (acessions MT353681-MT353684).

#### Electrophysiology

ND7/23 cell culture (ECACC 92090903, Sigma-Aldrich, Munich, Germany) and electrophysiological experiments were performed as described elsewhere ^31,32^. In detail, cells were cultured in Dulbecco’s Modified Eagle Medium (DMEM) supplemented with 5% (v/v) fetal bovine serum (FBS) and 1 µg/ml penicillin/streptomycin at 37 °C and 5% CO_2_. For experiments, cells were seeded on poly-D-lysine-coated coverslips at a density of 0.5–1.0×10^5^ cells/ml and supplemented with 1 µM all-*trans* retinal. The next day, cells were transiently transfected with 2 µg DNA using FuGENE^®^ HD (Promega, Madison, WI). Whole-cell patch-clamp recordings were performed at room temperature, two days after transfection with a 140 mM NaCl agar bridge as reference electrode and at membrane resistances ≥0.5 GΩ with an access resistance <10 MΩ. Patch pipettes were pulled to resistances of 1.5–2.5 MΩ using a P-1000 micropipette puller (Sutter, Novato, CA) and fire-polished. An AxoPatch200B and a DigiData400 were used to amplify and digitize signals, respectively. Signals were acquired using Clampex 10.4 (all from Molecular Devices, Sunnyvale, CA). A Polychrome V (TILL Photonics, Planegg, Germany) with the bandwidth set to 7 nm served as light source. The light was collimated into an Axiovert 100 microscope (Carl Zeiss, Jena, Germany) and controlled using a programmable shutter system (VS25 and VCM-D1; Vincent Associates, Rochester, NY). Buffer osmolarity was measured (Osmomat 3000basic, Gonotec, Berlin, Germany) and set to 320 mOsm for extracellular buffers and 290 mOsm for intracellular buffers using glucose. The pH was adjusted using N-methyl-D-glucamine or citric acid. Liquid junction potentials were calculated using Clampex 10.4 and corrected on-line. Extracellular buffers (Suppl. Table S1) were exchanged in random order to determine the ion selectivity, by manually adding and removing at least 4×1 ml of the respective buffer to the measuring chamber (volume ∼0.5 ml). Photocurrents were induced with 470-nm (viral ChRs) or 510-nm (CCMP277-1 and CCMP2087) light for 500 ms and recorded while the membrane potential was held at −80 to +40 mV in steps of 20 mV. Action spectra were recorded at -60 mV using 10-ms pulses of low-intensity light between 390 and 680 nm in steps of 10 nm. A motorized neutral-density filter wheel (Newport, Irvine, CA) was moved into the light path and controlled by a custom software written in LabVIEW (National Instruments, Austin, TX), to maintain an equal photon irradiance at all wavelengths. The wavelength sensitivity (*λ*_max_) was determined by applying a three-parametric Weibull function to the data normalized to the maximum photocurrent between 410 and 680 nm.

#### Confocal microscopy

For confocal imaging, ND7/23 cells were cultured as described above and seeded at a density of 0.2×10^5^ cells/ml in polymer-bottom, 35 mm µ-dishes (ibidi). Two days after transfection with 2 to 2.5 µg DNA using Fugene HD (Promega), confocal images were acquired using an FV1000 confocal laser scanning microscope (Olympus, Shinjuku, Tokyo, Japan) equipped with a 60x water immersion objective with a numerical aperture of 1.2 (UPlanSApo, Olympus). Protein localization was detected by exciting mCherry or eYFP with a 559 nm diode laser and a 515 nm argon laser, respectively (5% transmissivity for both). Acquired z-stacks were analyzed with ImageJ ^33^. Relevant z-planes were z-projected for representative images of membrane fluorescence.

## Supplementary Figures

**Fig. S1.**
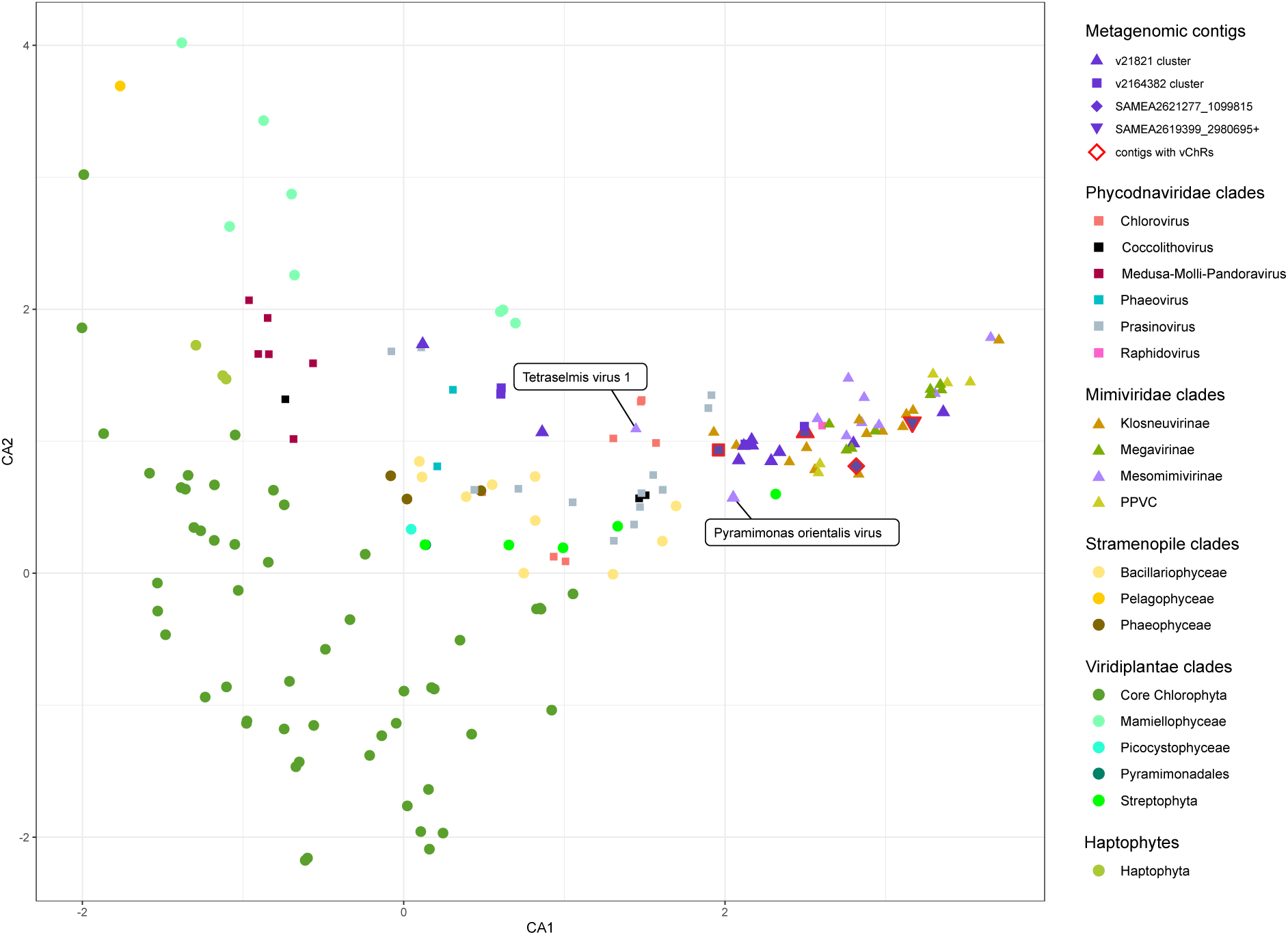
Ordination of the metagenomic contigs containing putative viral ChRs and related contigs, genomes of viruses from the *Mimiviridae* and *Phycodnaviridae* and genomes of several photosynthetic organisms in the canonical axes of the correspondence analysis of tetranucleotide frequencies.

**Fig. S2.**
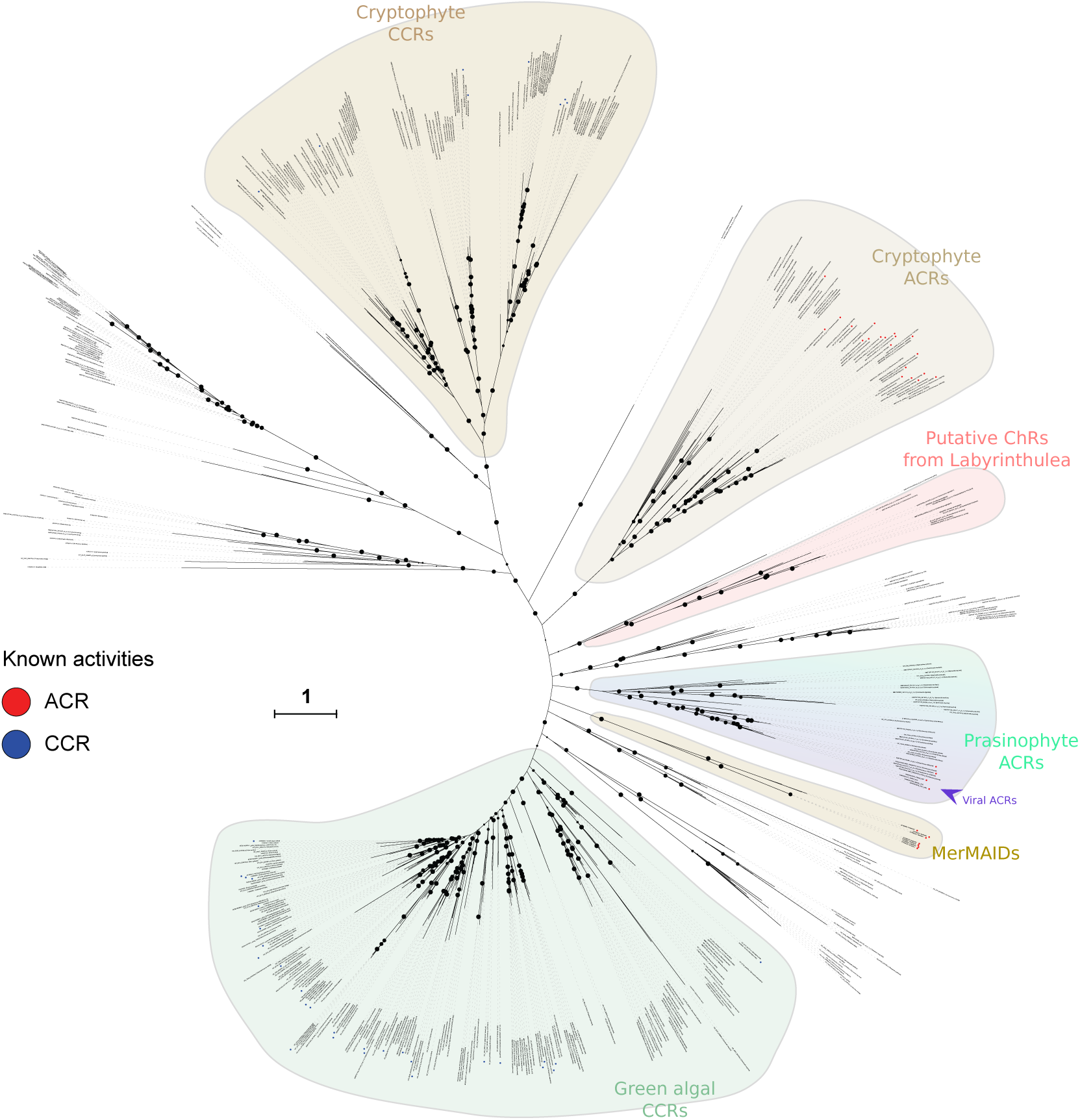
Phylogenetic analysis of known and putative channelrhodopsins. The prasinophyte and viral ACRs form a well-supported clade not nested in any of the described families of ChRs. The ultrafast bootstrap support values are indicated by circles (70-100 range). The sequences and the phylogenetic tree are available in Suppl. File 3a,b.

**Fig. S3.**
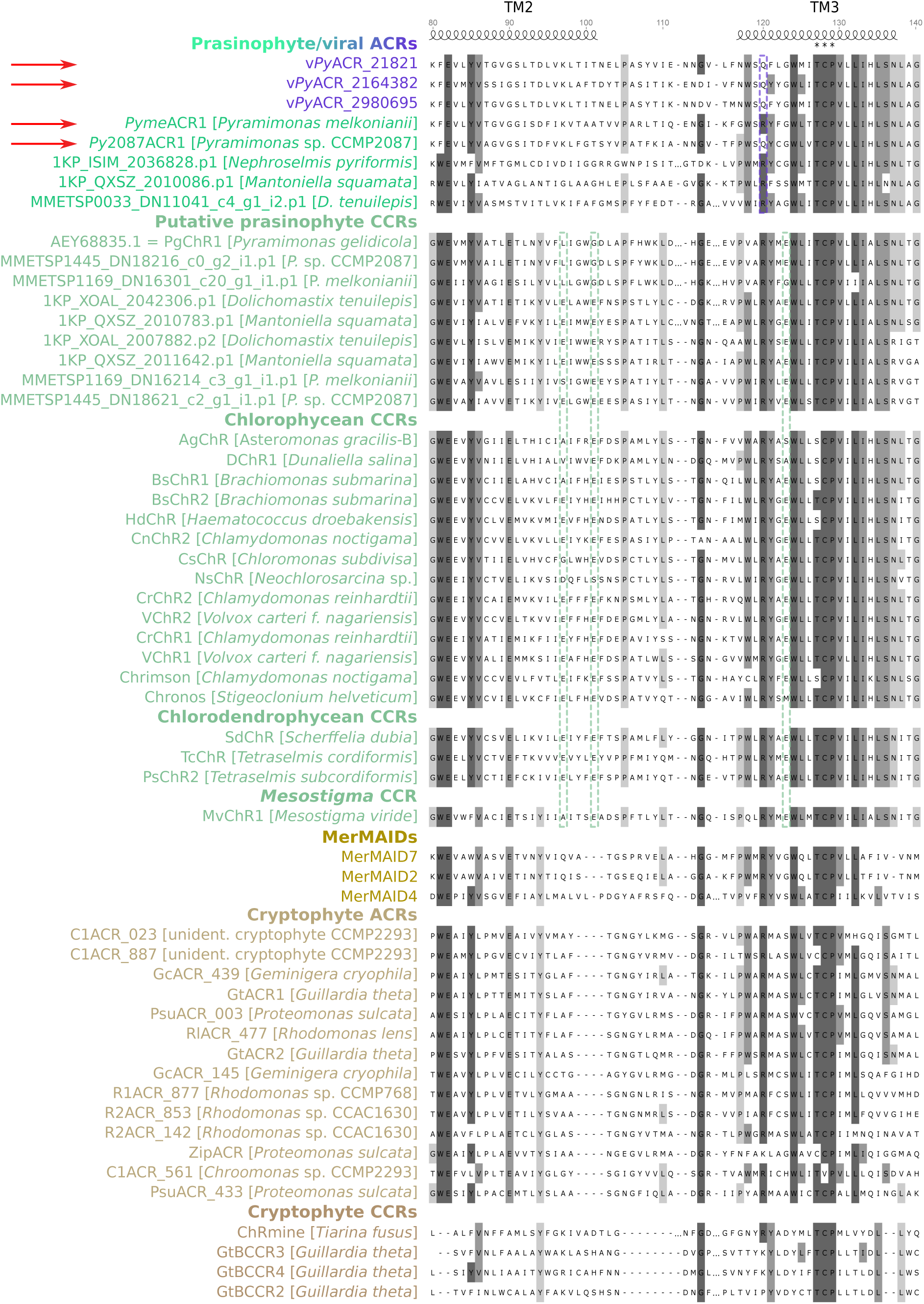
Alignment of the transmembrane domains 2 and 3 (TM2 and TM3) of different channelrhodopsins. Representative sequences of the ChRs with known activities were obtained by clustering at 60% identity level. The prasinophyte ChRs tested in this study are marked with red arrows, the activity of the other proteins from this group is putative. Green algal CCRs are subdivided into taxonomic groups (cf. Fig. S2). The location of the TMs and the position numeration correspond to the structure of *Cr*ChR2 (PDB: 6EID); the alignment positions are highlighted according to conservation level; the Asp motifs in green algal CCRs are marked with dashed green frames; the XCP motif in TM3 is marked with asterisks; the unique Arg>Gln substitution in TM3 in viral ACRs and *Py*2087ACR1 is indicated with a dashed lilac frame.

**Fig. S4.**
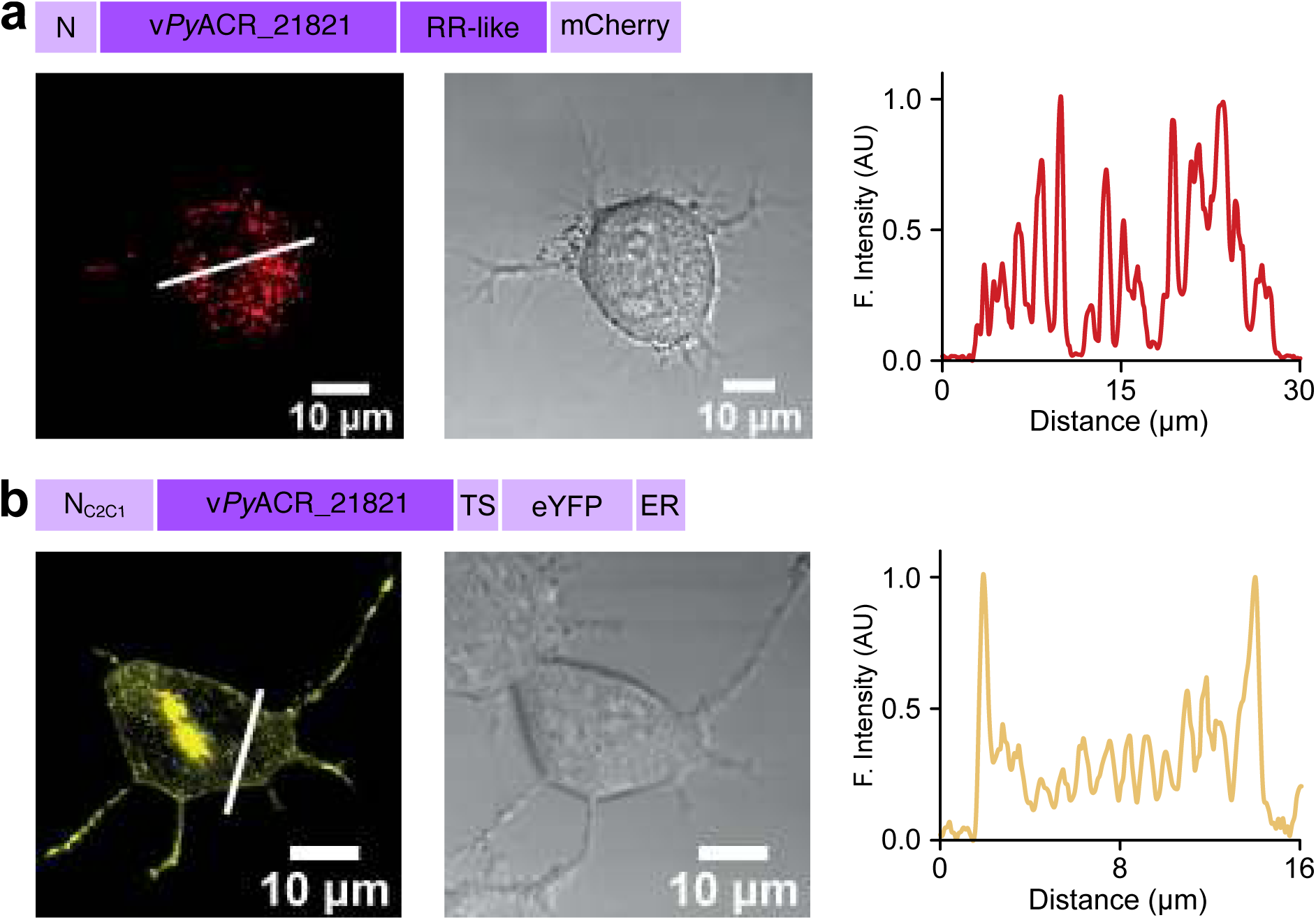
Membrane targeting of v*Py*ACR_21821. Confocal images of ND7/23 cells after two days’ expression of full-length (a) or membrane-targeted (b) v*Py*ACR_21821. Fluorescence (left) of mCherry is shown in red and of eYFP in yellow. Fluorescence intensity profiles on the right were measured at the locations indicated by a thin white line in the fluorescence images.

**Fig. S5.**
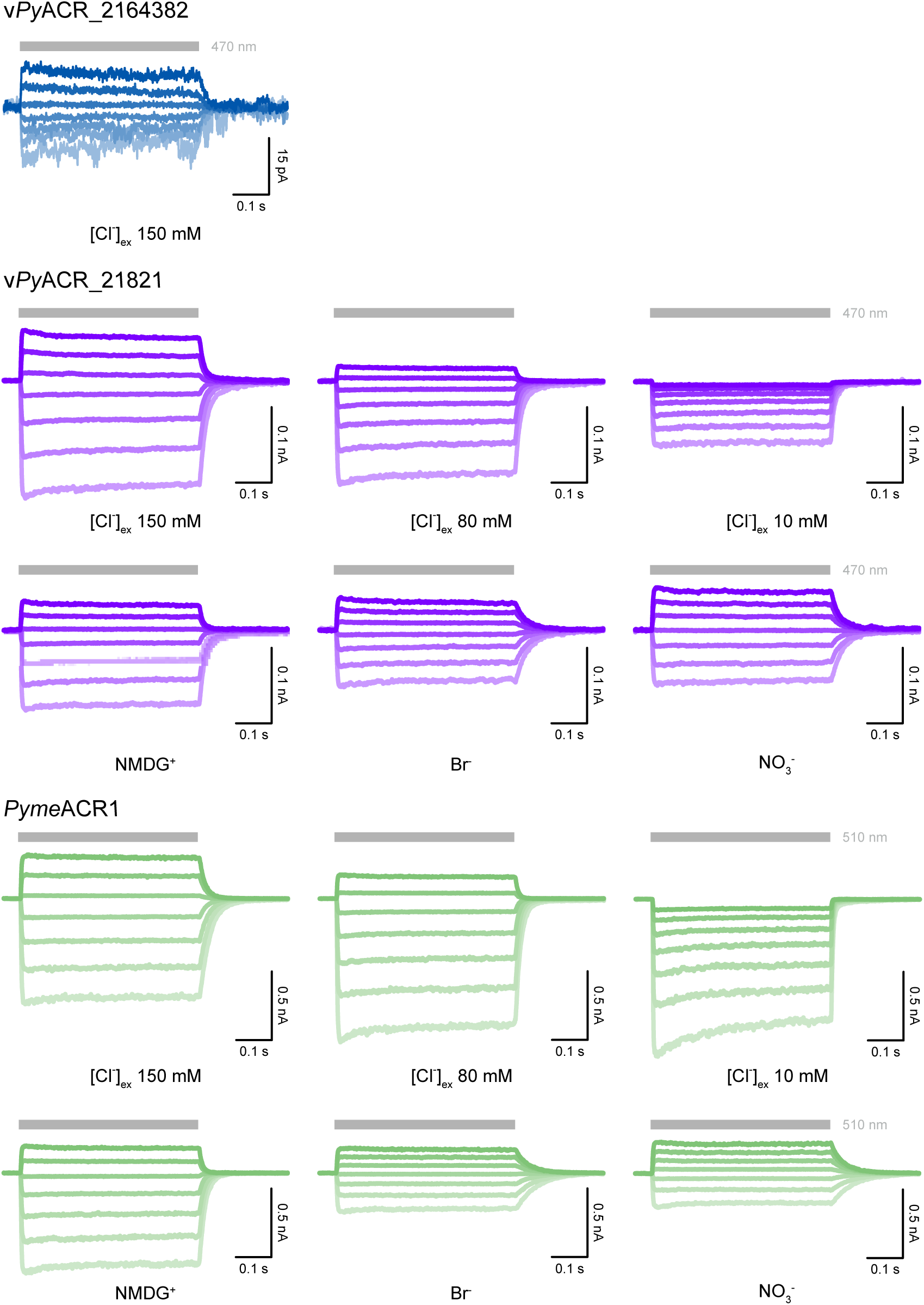
Photocurrent traces of v*Py*ACR_2164382, v*Py*ACR_21821, and *Pyme*ACR1. Photocurrents were induced with light of indicated wavelengths (gray bars) and recorded at membrane potentials between −80 mV (lightest colored lines) and +40 mV (darkest colored lines) in steps of 20 mV. The external buffer contained high concentrations of the indicated ions (see Suppl. Table 1 for buffer compositions).

**Fig. S6.**
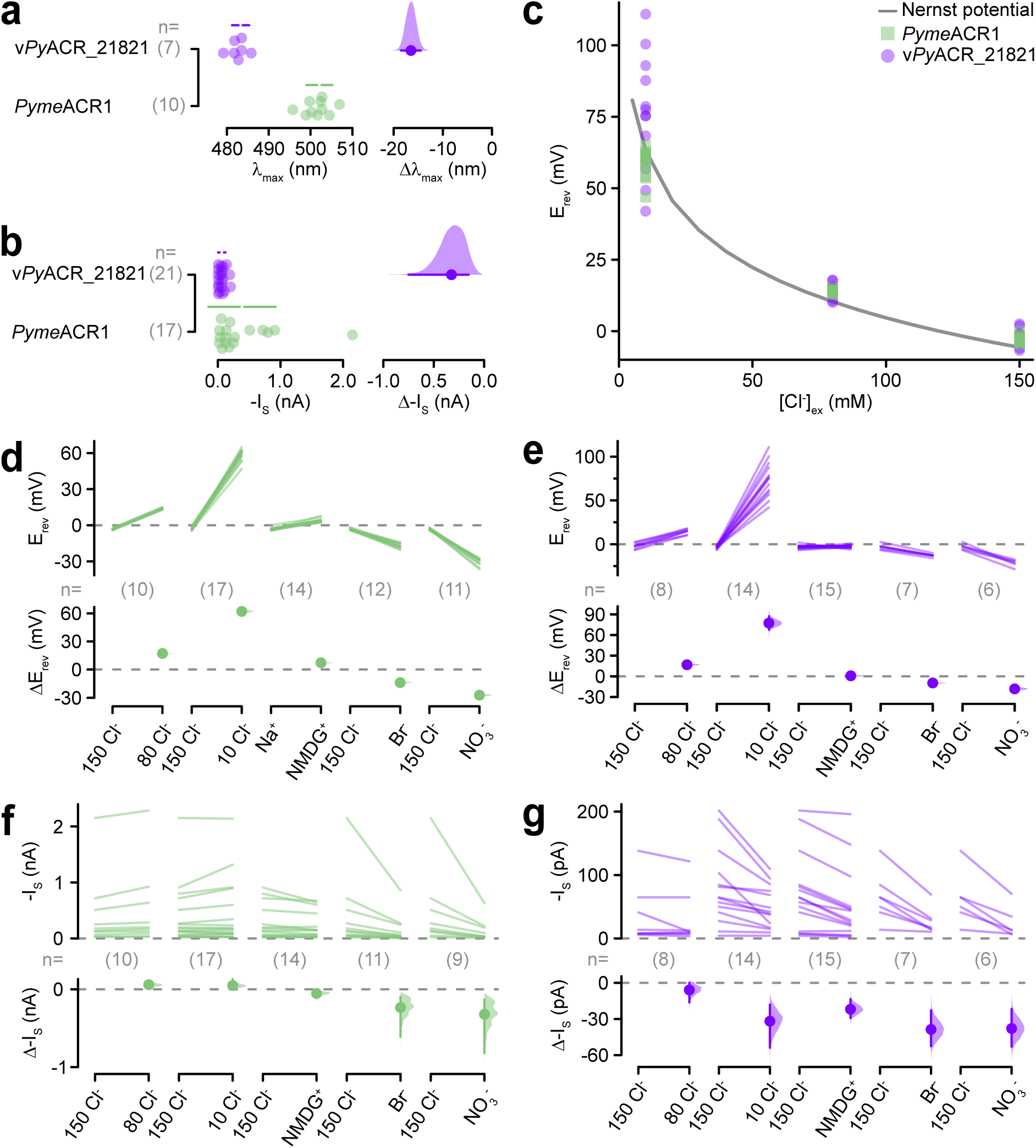
Theoretical vs experimental reversal potential and estimation plots for electrophysiological data. **(a) & (b)** Estimation plots of *λ*_max_ for v*Py*ACR_21821 and *Pyme*ACR1 (a) and stationary photocurrent amplitudes (I_S_) at -60 mV (b). **(c)** Comparison of the theoretical Nernst potential for chloride (line) with the experimentally derived reversal potentials (E_rev_) for v*Py*ACR_21821 (purple) and *Pyme*ACR1 (green) at external chloride concentrations ([Cl^-^]_ex_) of 10 mM, 80 mM, and 150 mM. **(d, e)** Estimation plots of paired E_rev_ (top) and resulting ΔE_rev_ (bottom) upon exchange of the external buffer as indicated (see Suppl. Table S1 for buffer compositions) for *Pyme*ACR1 (d) and v*Py*ACR_21821 (e). **(f, g)** Estimation plots of paired stationary photocurrent amplitudes at -60 mV (-I_S_) and resulting shifts (Δ-I_S_) upon exchange of the external buffer as indicated (see Suppl. Table S1 for buffer compositions) for *Pyme*ACR1 (f) and v*Py*ACR_21821 (g). The estimation plots show the mean difference between test and control group. Both groups are plotted as pairs or as single data points (a and b) with the mean values (white dot) ± standard deviation (solid line). The mean difference is indicated by a solid dot with each bootstrap sampling distribution indicated by a filled curve with the error bars in indicating the 95% confidence interval. The number of biological replicates is indicated (n). Source data are provided Suppl. File 5.

**Fig. S7.**
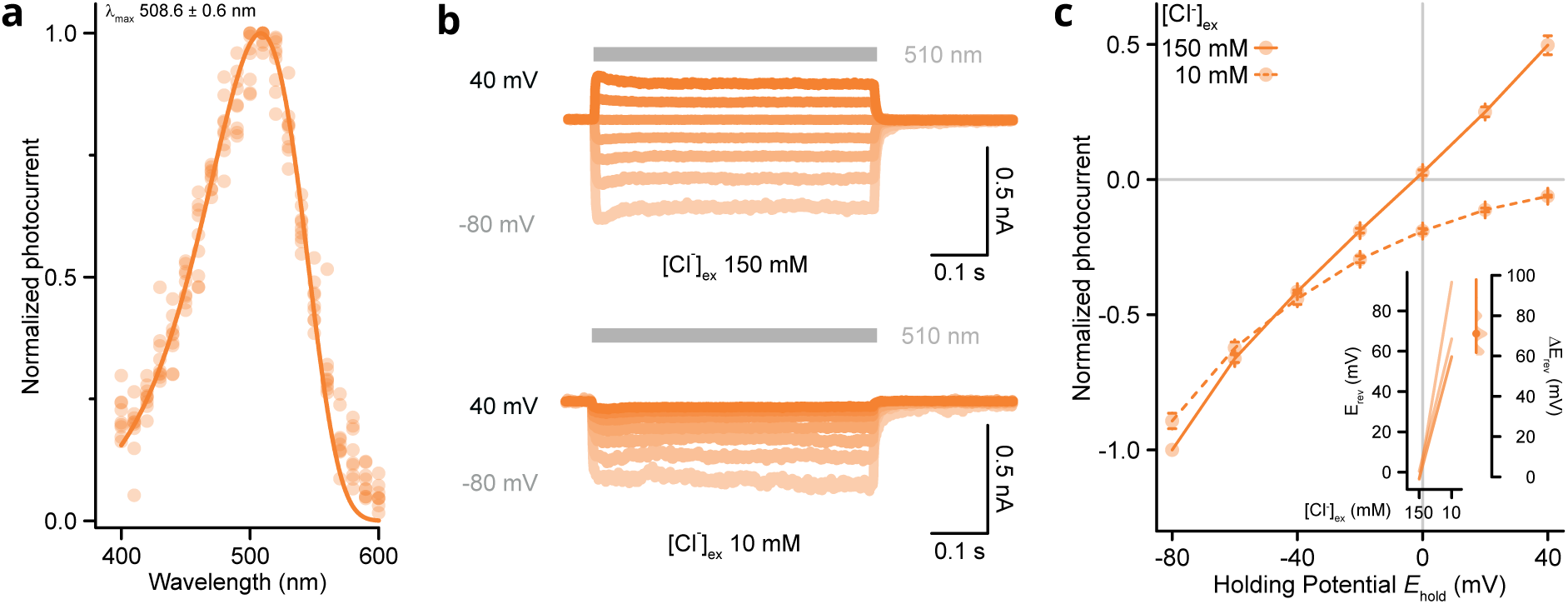
Basic electrophysiological characterization of *Py*2087ACR1. **a**, Action spectrum normalized to the maximum photocurrent. Dots are single measurements (n = 9) fitted (solid line) to determine maximum activity wavelength (*λ*_max_) indicated as mean ± SEM. **b**, Example current traces recorded with an external chloride concentration ([Cl^-^]_ex_) of 150 mM or 10 mM (internal [Cl^-^]: 120 mM) at membrane potentials between −80 mV and +40 mV in steps of 20 mV. Photocurrents were elicited with 510 nm light (gray bars). c, Current-voltage relationship of photocurrents at [Cl^-^]_ex_ of 150 mM (solid line; n = 4) and 10 mM (dashed line; n = 4). Inset shows a paired estimation plot of the determined reversal potentials (E_rev_) for the respective measurements (left) and the resulting reversal potential shifts (ΔE_rev_; right). The mean is indicated by a solid dot with the error bars indicating 95% confidence interval. The bootstrap sampling distribution is indicated by a filled curve. Source data are provided in Suppl. File 5.

**Figure S8.**
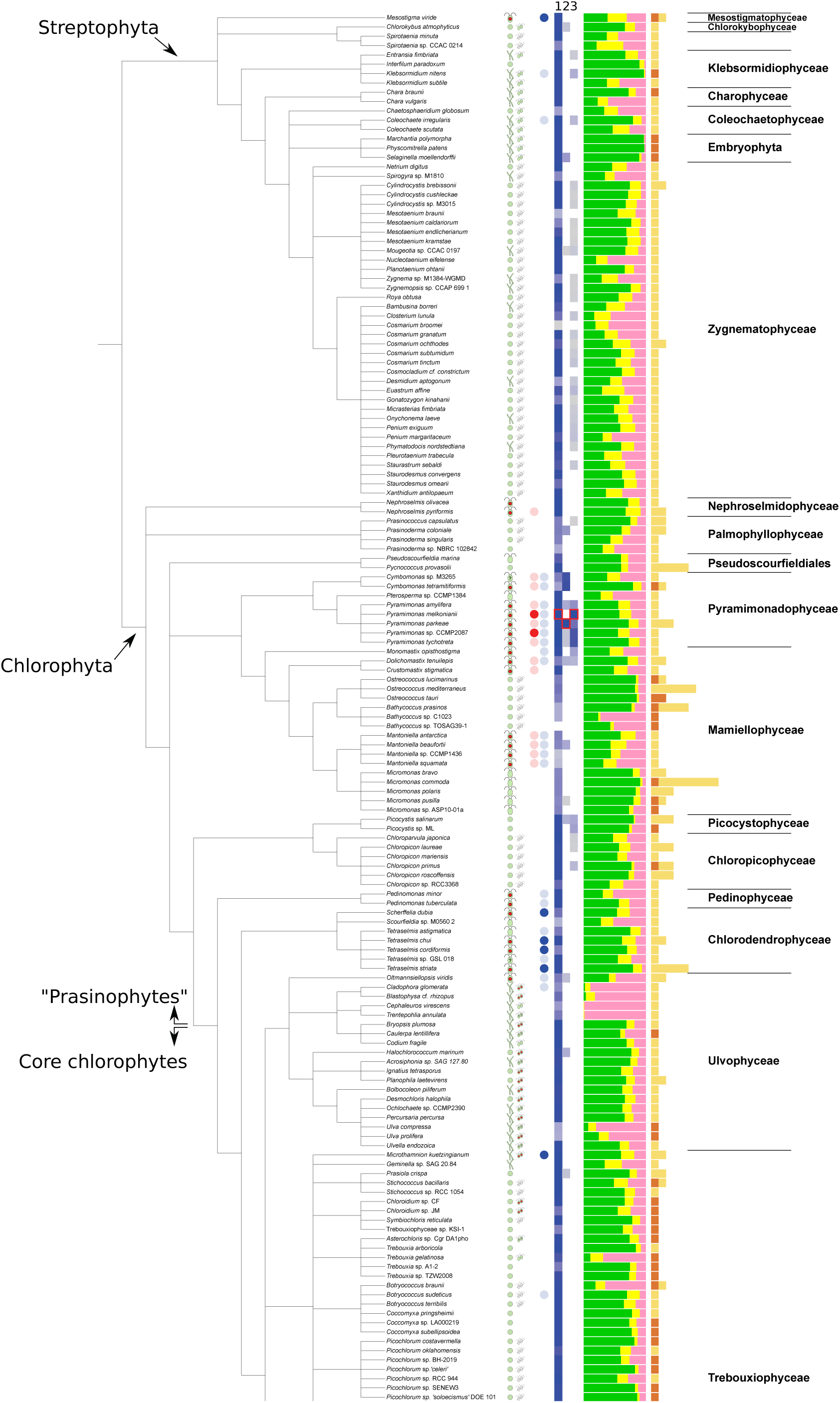
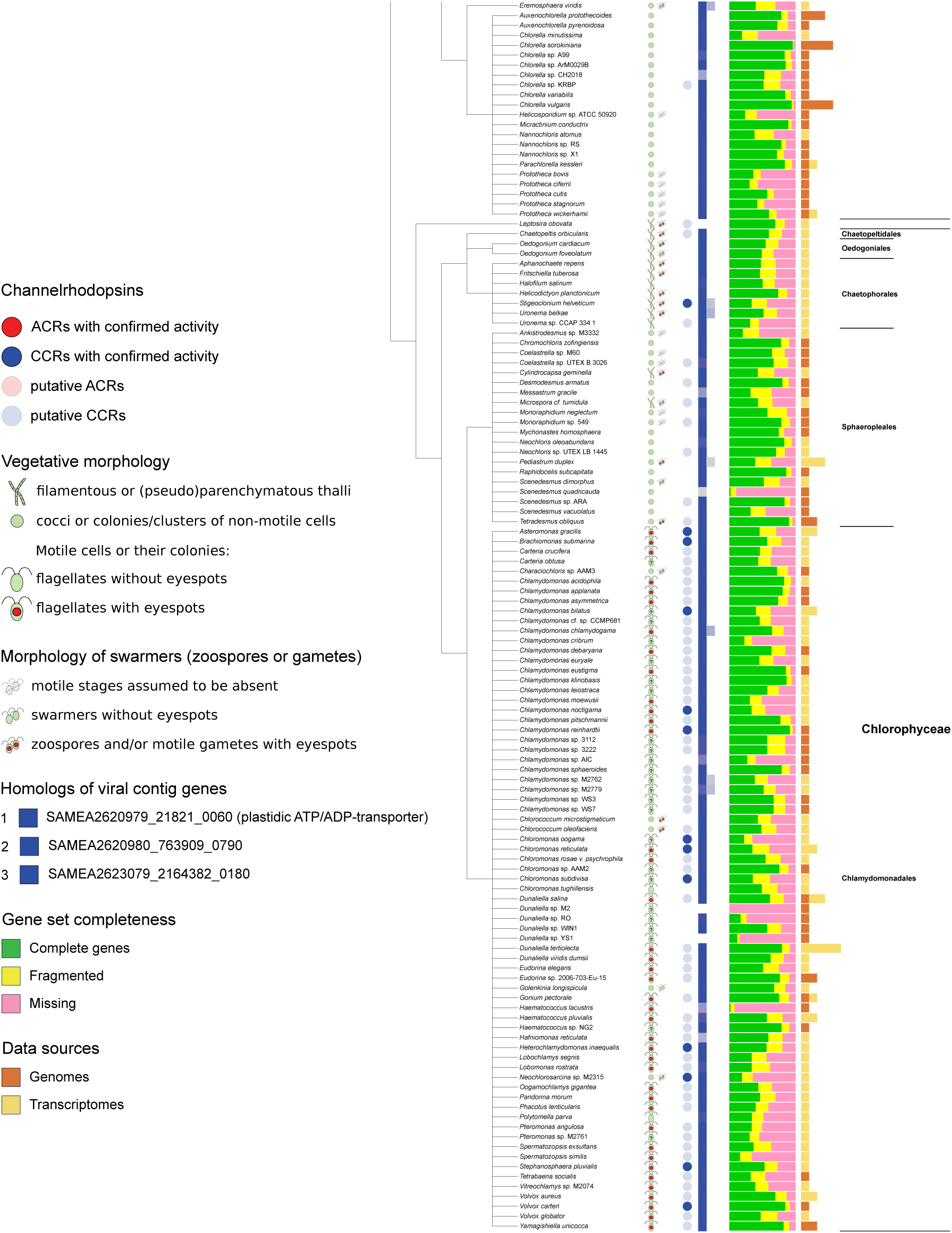
Distribution of confirmed and putative channelrhodopsins among green algae. The cladogram reflects the consensus topology of green algal phylogeny largely based on Leliaert et al ^17^ and further refined based on ^13,34^. Available genome and transcriptome assemblies were merged together on the level of the species (see Suppl. File 4).

**Fig. S9.**
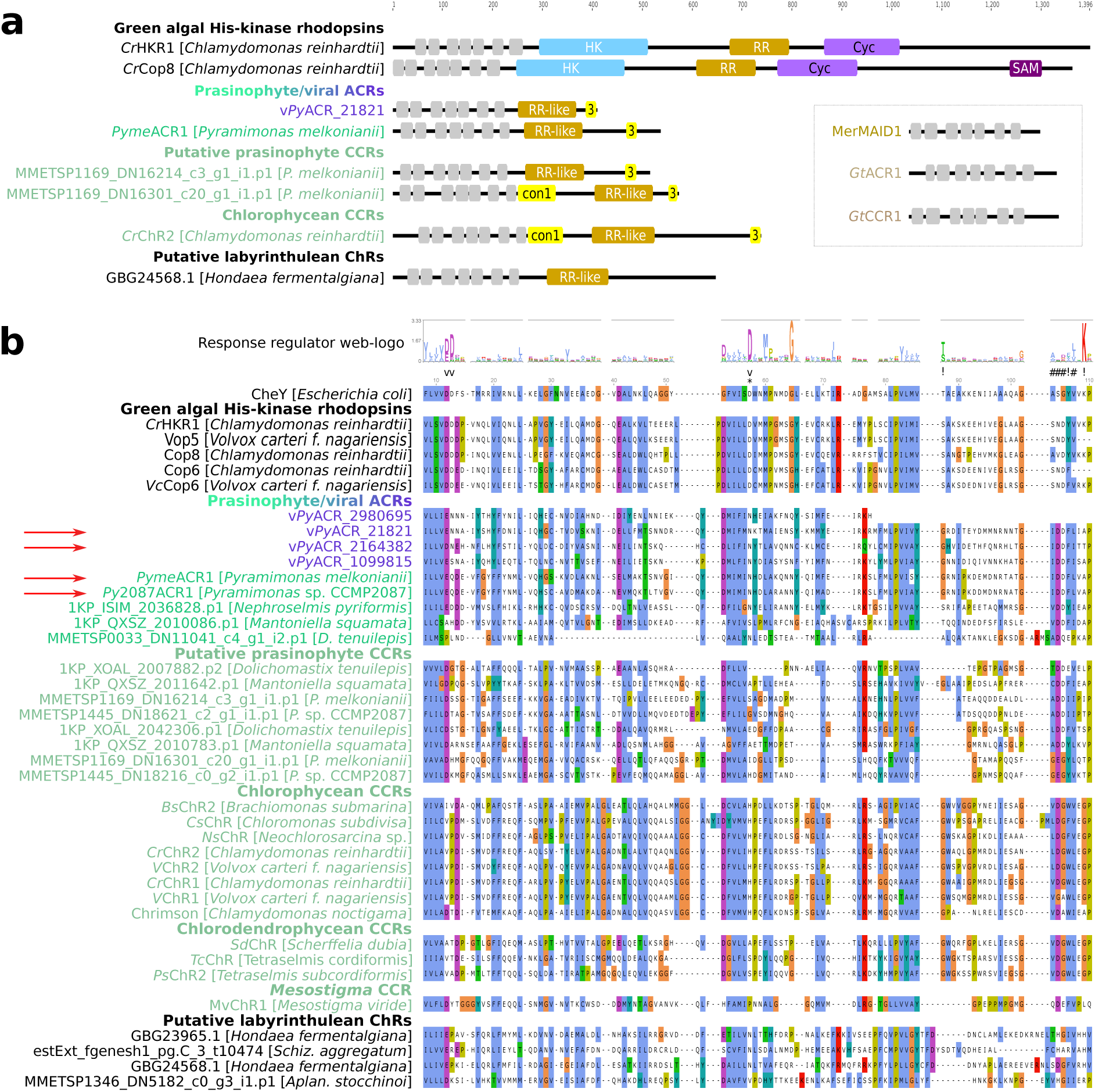
Domain organization in different channelrhodopsin families, compared to the related family of green algal His-kinase rhodopsins (HKRs). **(a)** Domains in the intracellular C-terminal extensions in representative ChRs and HKRs. Prasinophyte and chlorophycean CCRs belong to green algal CCRs. The inset shows the three ChR families without C-terminal extensions. Gray boxes indicate rhodopsin TM domains. HK — His-kinase domains, RR(-like) — response regulator(-like) domains, Cyc — nucleotide cyclase domains, SAM — sterile alpha motif domain, con1 and 3 — conserved regions 1 and 3 first described CCRs from *Chlamydomonas* and *Volvox* ^22^. **(b)** Response regulator-like domains in three families of channelrhodopsins compared to their functional homologs: CheY and response regulator domains from green algal His-kinase rhodopsins. Green algal CCRs are subdivided into taxonomic groups. The numeration corresponds to residue positions in CheY; above the alignment is a web-logo for the seed alignment of the Pfam profile Response_reg; response regulator active sites are indicated with symbols (as summarized in NCBI CDD: cd00156): * — phosphorylation site, v — divalent cation binding sites, # — dimerization interface, ! — other active sites.

**Figure S10.**
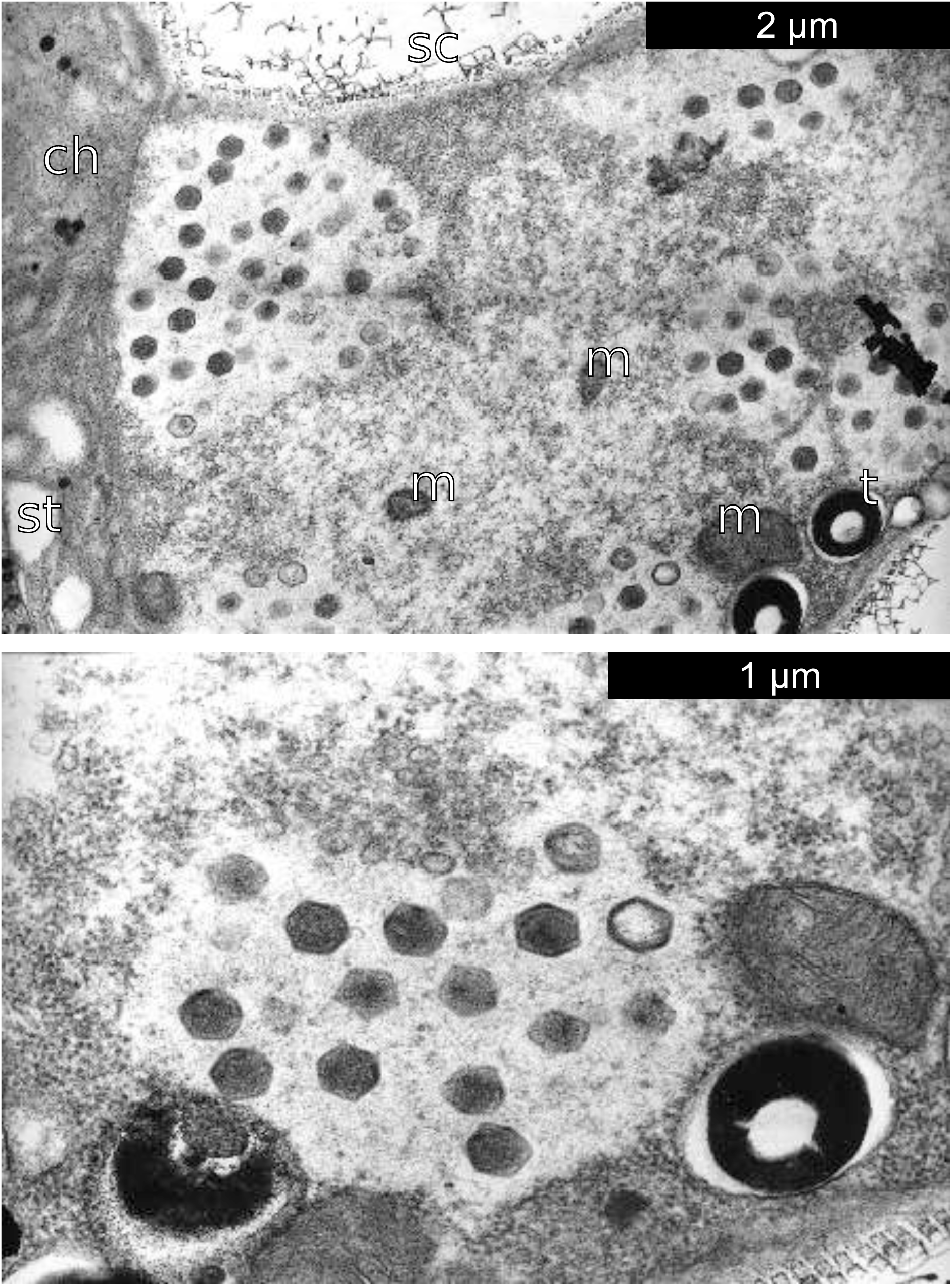
TEM micrograph of a *Pyramimonas pseudoparkeae* cell infected with a giant virus. Upper panel: general overview of viral factories and surrounding cellular structures, lower panel: closeup of a viral factory. ch — chloroplast, m — mitochondria, t — trichocyst, st — starch grains, sc — scales. The diameter of the mature viral particles is ca. 199 nm. Courtesy of Richard Pienaar.

**Figure S11.**
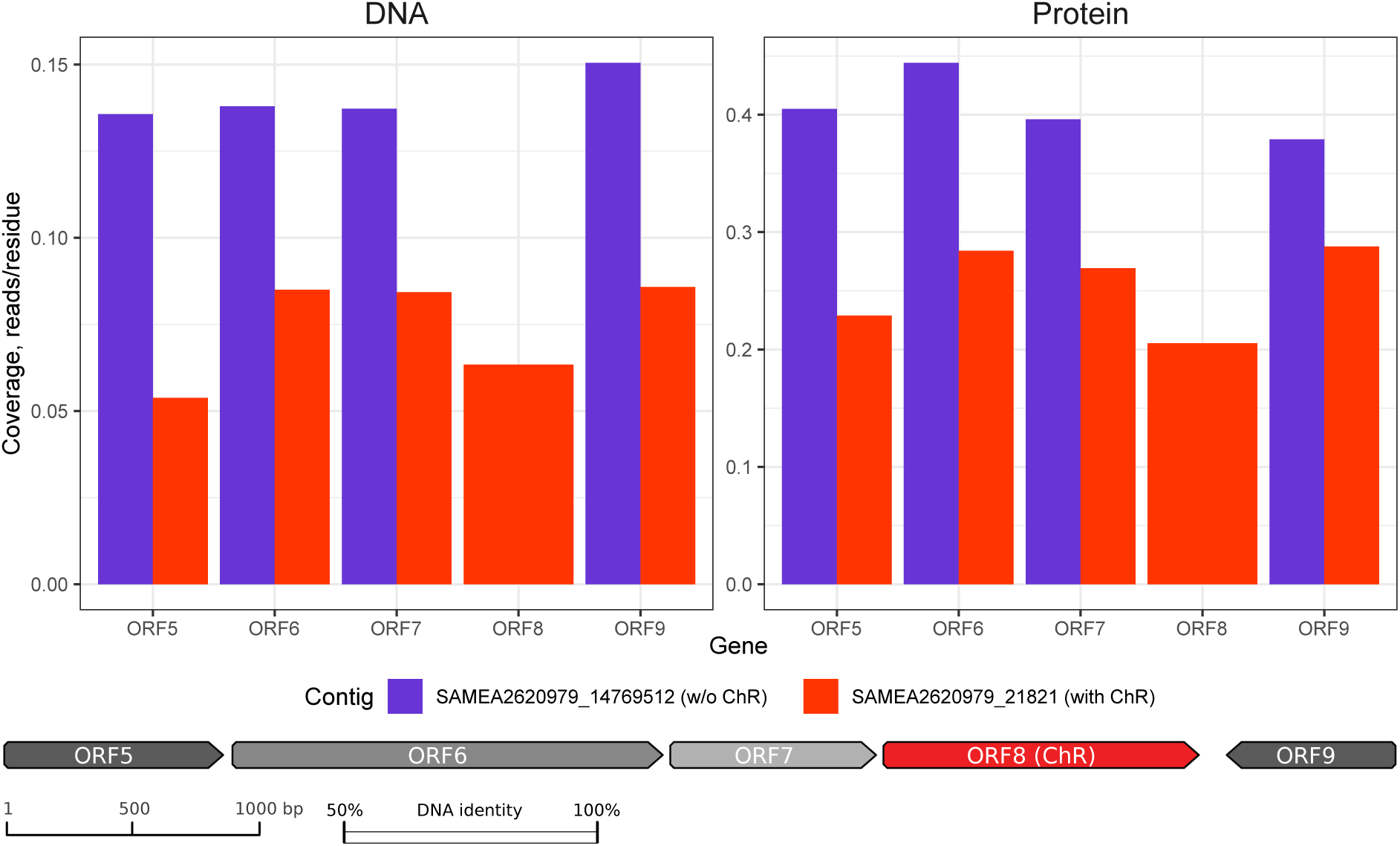
Abundance of the two variants of the genomic location around the channelrhodopsin gene at station TARA_067.

**Figure S12.**
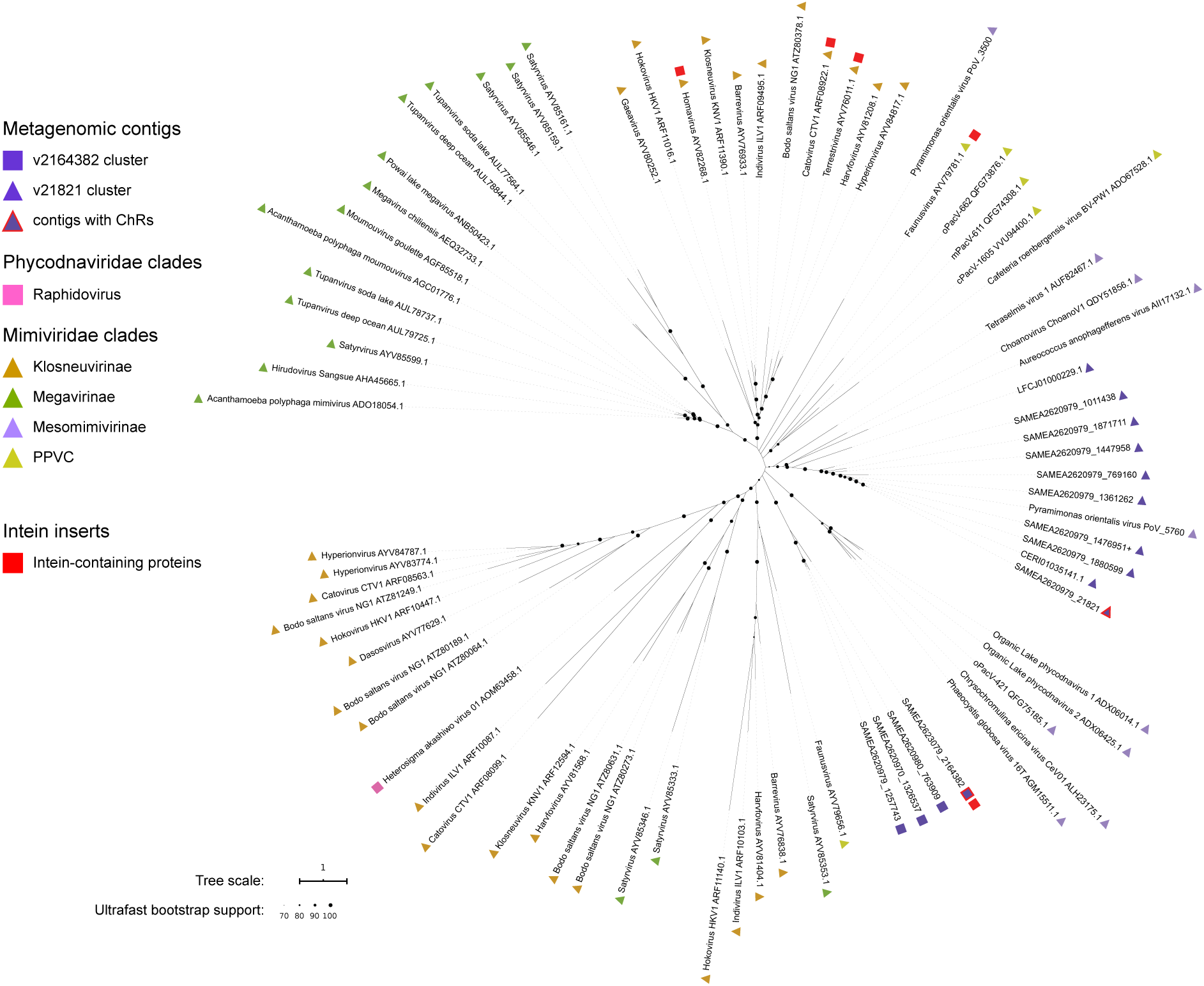
D5-like helicase-primase phylogeny. Proteins with intein inserts are indicated with red squares.

**Fig. S13.**
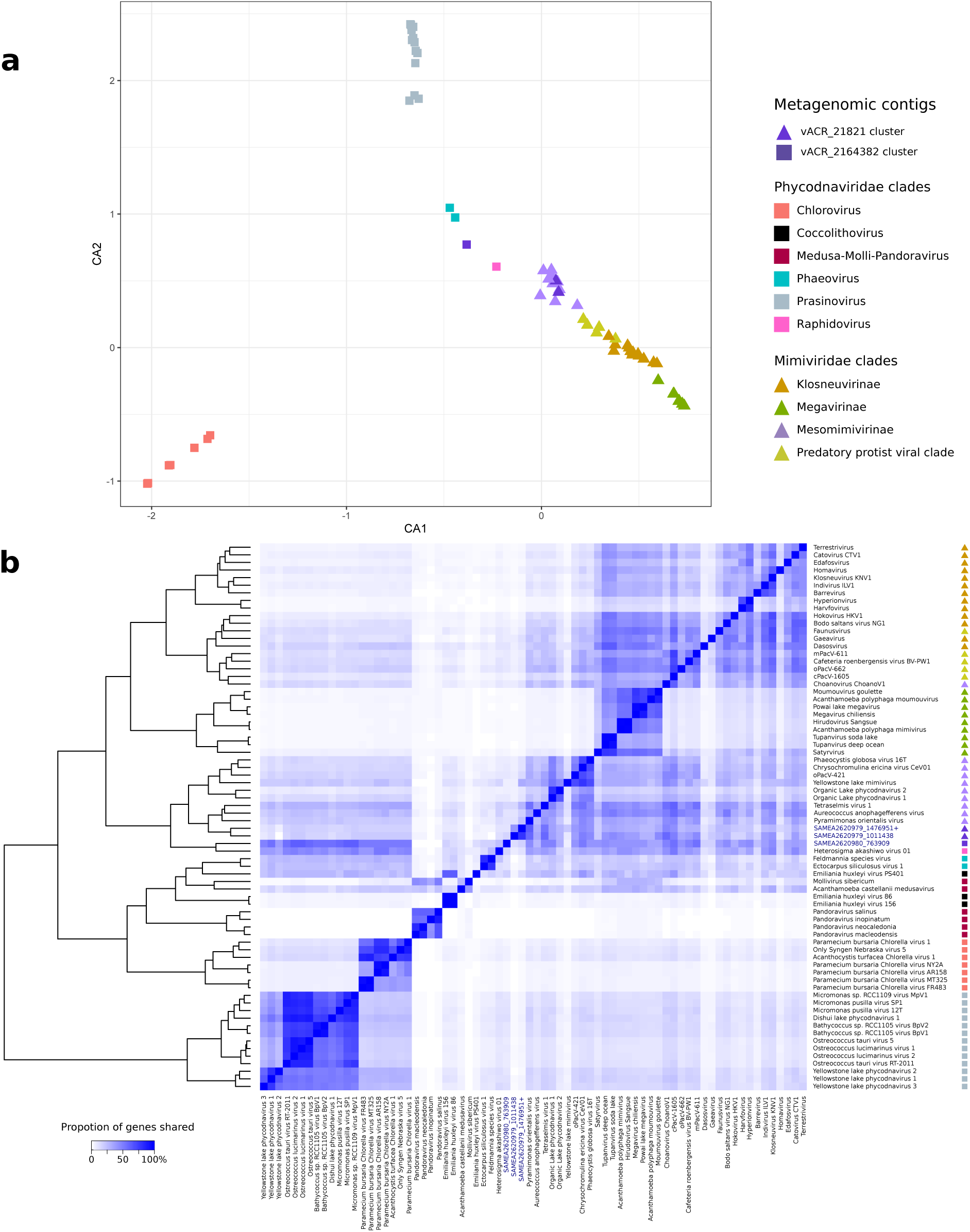
Gene sharing analyses for the members of the *Phycodna-* and *Mimiviridae* and the metagenomic contigs. **(a)** Ordination of long metagenomic contigs and genomes of related viruses in the canonical axes of correspondence analysis based on orthogroup presence/absence. The genomes of *Coccolithovirus, Medusavirus, Mollivirus* and *Pandoravirus* appeared as outliers with respect to the core *Phycodnaviridae* and the *Mimiviridae* and were omitted in the second round of this analysis as presented here. **(b)** Heatmap and clustering of long metagenomic contigs and viruses from the *Phycodnaviridae* and *Mimiviridae* families based on orthogroup sharing. Color intensity reflects the proportion of genes shared between a genome pair (relative to the genome in the row).

**Fig. S14.**
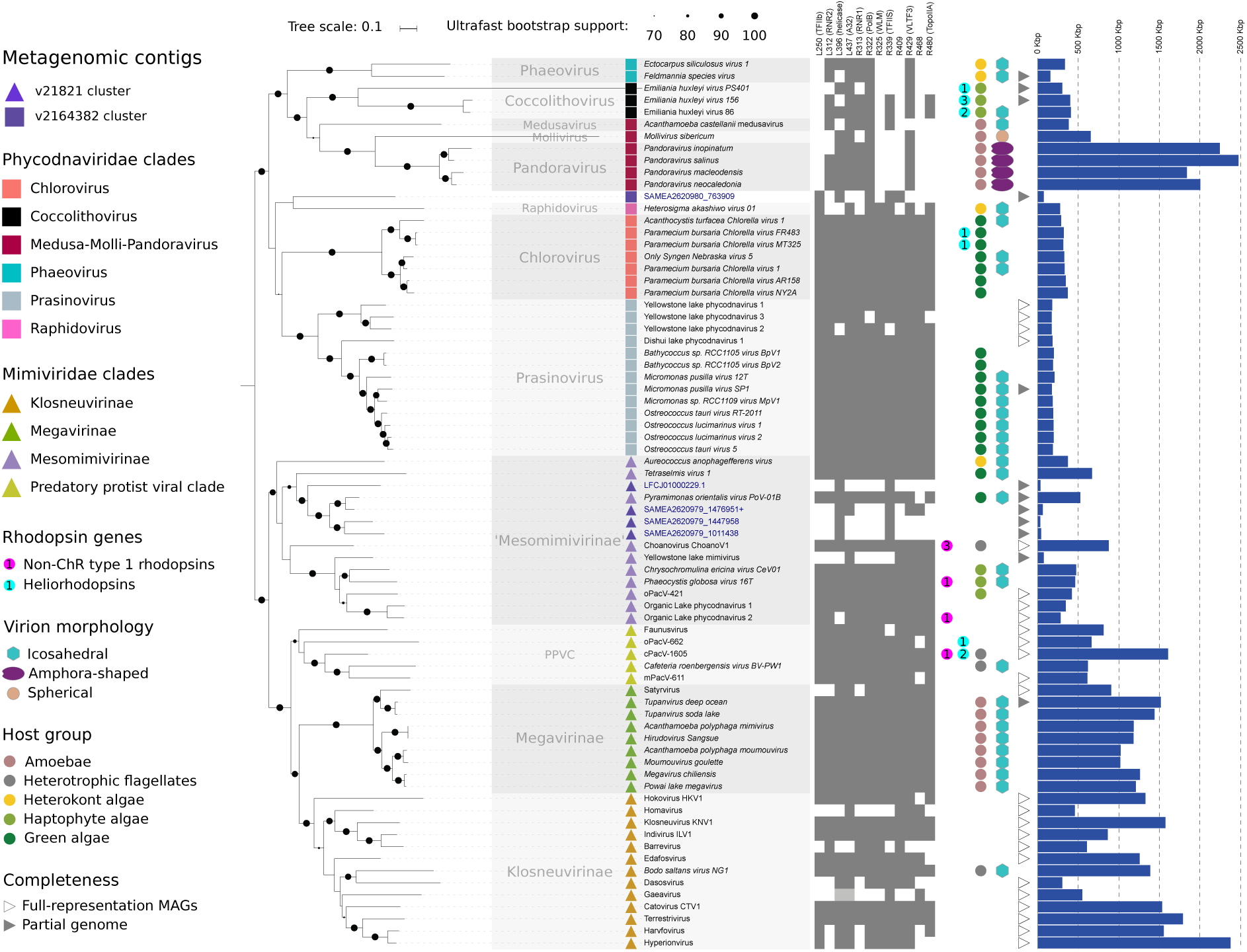
Phylogenetic relationships among *Mimi-* and *Phycodnaviridae*, and the lineages including ChR-containing viruses, showing the distribution of rhodopsin genes, virion morphology, known host groups and genome sizes. The relationships between the long metagenomic contigs and their shorter relatives containing ChRs are shown in Fig. 1c. The phylogenetic reconstruction is based on 12 near-universal genes and their names in *Mimivirus* (with functions in parentheses) and distribution in the alignment are depicted. The protein sequences for each of the 12 genes are available in Suppl. File 6a,b.

**Fig. S15.**
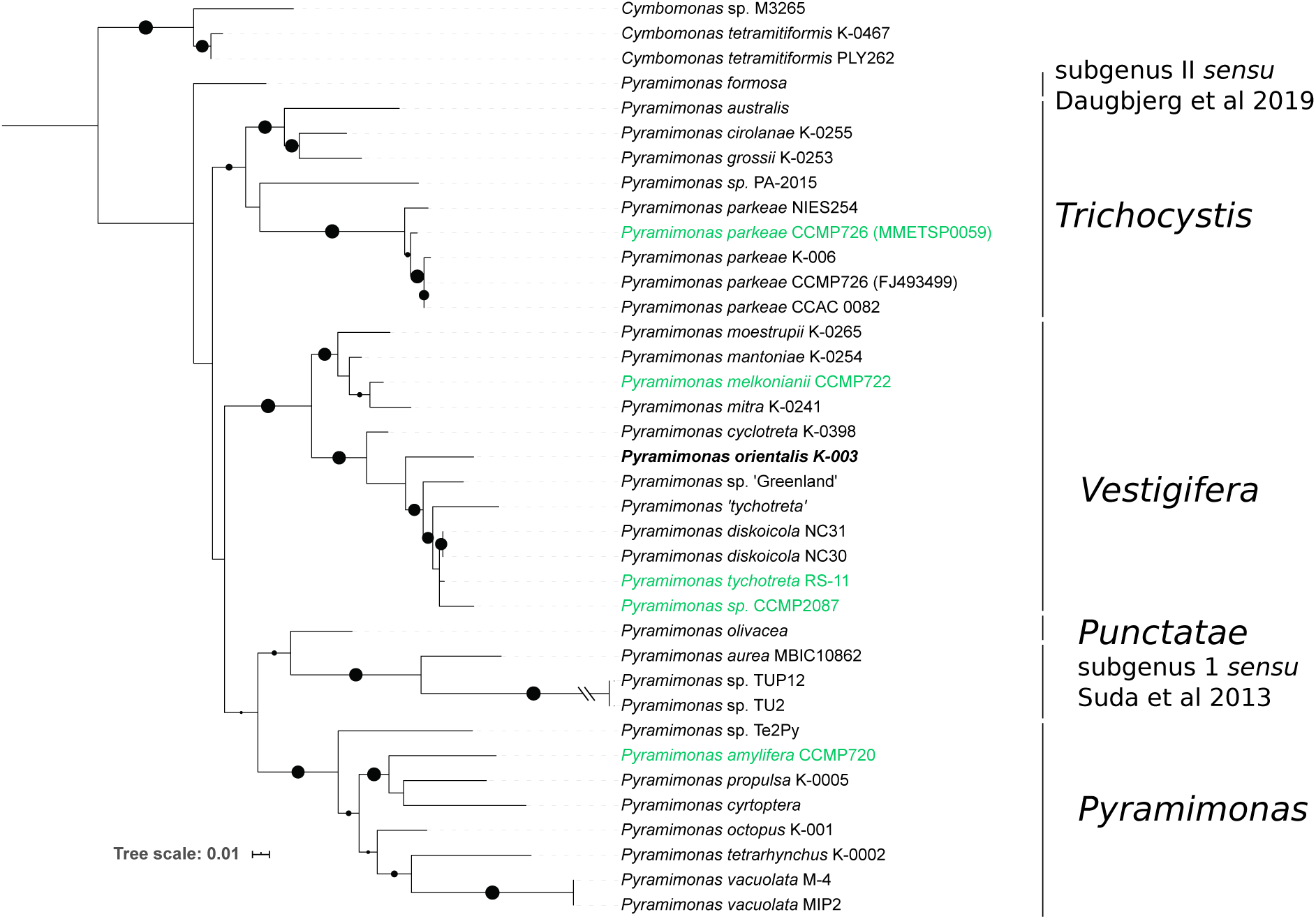
Nucleotide chloroplast rbcL phylogeny of *Pyramimonas*. Strains appearing in Fig. 1c are indicated in green and the host of PoV-01B is indicated in bold. The clade names (subgenera) follow the previous studies ^34–36^. Dots indicate ultrafast bootstrap support values (70-100 range), the tree is rooted using rbcL sequences from *Cymbomonas*. rbcL sequence alignment is available in Suppl. File 7.

## Supplementary Tables

**Supplementary Table S1.** Composition of intra- and extracellular buffers for electrophysiological experiments. All concentrations are given in mM, LJPs are listed in mV. Asp, aspartate; EGTA, ethylene glycol tetraacetic acid; HEPES, 4-(2-hydroxyethyl)-1-piperazineethanesulfonic acid; LJP, liquid junction potential; NMDG, N-Methyl-D-glucamine

|  | NaCl | KCl | MgCl<sub>2</sub> | CaCl<sub>2</sub> | CsCl | NaAsp | NMDG | HCl | NaBr | NaNO<sub>3</sub> | HEPES | EGTA | LJP |
| --- | --- | --- | --- | --- | --- | --- | --- | --- | --- | --- | --- | --- | --- |
| *Intra* |  |  |  |  |  |  |  |  |  |  |  |  |  |
| Std. | 110 | 1 | 2 | 2 | 1 | - | - | - | - | - | 10 | 10 | - |
| *Extra* |  |  |  |  |  |  |  |  |  |  |  |  |  |
| 150 Cl<sup>-</sup> | 140 | 1 | 2 | 2 | 1 | - | - | - | - | - | 10 | - | 0.6 |
| 80 Cl<sup>-</sup> | 70 | 1 | 2 | 2 | 1 | 70 | - | - | - | - | 10 | - |  |
| 10 Cl<sup>-</sup> | - | 1 | 2 | 2 | 1 | 140 | - | - | - | - | 10 | - | -12.6 |
| NMDG<sup>+</sup> | 1 | 1 | 2 | 2 | 1 | - | 140 | 140 | - | - | 10 | - | 6.3 |
| Br<sup>-</sup> | - | 1 | 2 | 2 | 1 | - | - | - | 140 | - | 10 | - | 1.0 |
| NO<sub>3</sub><sup>-</sup> | - | 1 | 2 | 2 | 1 | - | - | - | - | 140 | 10 | - | -0.3 |

## List of supplementary data files

**Suppl. File 1**. Annotated metagenomic contigs analyzed in this study including the contigs containing ChR genes, file in Genbank format.

**Suppl. File 2.** Annotated genome draft of *Pyramimonas orientalis virus PoV-01B*, file in Genbank format.

**Suppl. File 3a.** Database of collected ChR sequences, xlsx file: non-redundant protein sequences, confirmed activities, representative sequences after 98% identity clustering, representative sequences after 100% identity clustering of trimmed rhodopsin domains.

**Suppl. File 3b.** Final dataset of rhodopsin domains and the phylogenetic tree, nexus file.

**Suppl. File 4**. List of analyzed green algal transcriptomes and transcriptomes and viral genomes, xlsx file. Sheet A: Green algal species analyzed for the presence of ACRs and CCRs: taxonomy and data sources (strais: accessions). References are provided for data sources other than MMETSP ^10^ and 1KP ^11^. Original species identifications are provided in parenthesis when different. Sheet B: Viral genomes used for clarification of the origin of the metagenomic contigs containing ChR genes.

**Suppl. File 5.** The source data from Ephys for electrophysiological experiments used in Figs. 2, S6 and S7, xlsx file.

**Suppl. File 6a.** The twelve genes used for phylogenetic analysis of the *Mimiviridae* and *Phycodnaviridae*, xlsx file: L250 (TFIIb), L312 (RNR2), L396 (helicase), L437 (A32), R313 (RNR1), R322 (PolB), R325 (WLM), R339 (TFIIS), R409, R429 (VLTF3), R468, R480 (TopoIIA).

**Suppl. File 6b**. Phylogenetic tree of the *Mimiviridae* and *Phycodnaviridae* and the metagenomic contigs, nexus file.

**Suppl. File 7.** Alignment of rbcL gene sequences from *Pyramimonas* and Cymbomonas species and the corresponding phylogenetic tree, nexus file.

**Submitted to Genbank - Constructs.gbk.** Mammalian codon-optimized constructs of prasinophyte and viral ACR coding sequences used for expression. (Genbank accession numbers MT353681-MT353684).

